# Regulation of RAS palmitoyltransferases by accessory proteins and palmitoylation

**DOI:** 10.1101/2022.12.12.520165

**Authors:** Anlan Yang, Shengjie Liu, Yuqi Zhang, Jia Chen, Shan Feng, Jianping Wu, Qi Hu

## Abstract

**Summary:** Palmitoylation of cysteine residues at the C-terminal hypervariable regions in human HRAS and NRAS, which is necessary for the RAS signaling, is catalyzed by the acyltransferase DHHC9 in complex with its accessory protein GCP16. The molecular basis for the acyltransferase activity and the regulation of DHHC9 by GCP16 is not clear. Here we report the cryo-EM structures of the human DHHC9-GCP16 complex and its yeast counterpart — the Erf2-Erf4 complex, demonstrating that GCP16 and Erf4 are not directly involved in the catalytic process but stabilize the architecture of DHHC9 and Erf2, respectively. We found that a phospholipid binding to an arginine-rich region of DHHC9 and palmitoylation on three residues (C24, C25, and C288) were essential for the catalytic activity of the DHHC9-GCP16 complex. Furthermore, we showed that GCP16 also formed complexes with DHHC14 and 18 to catalyze RAS palmitoylation. These findings provide insights into the regulation mechanism of RAS palmitoyltransferases.

## Introduction

The RAS GTPases switch between the inactive GDP-bound state and the active GTP-bound state, and this process is regulated by guanine nucleotide exchange-factors (GEFs) and GTPase-activating proteins (GAPs) (Simanshu et al., 2017). In addition to the nucleotide exchange, lipidation of RAS adds another layer of regulation to the RAS signaling pathway. All RAS isoforms have a CAAX motif at their C-terminus in which the cysteine residue can be prenylated; after prenylation, the last three residues AAX is removed by RCE1 (RAS converting CAAX endopeptidase 1) and the exposed carboxyl group of the prenylated cysteine is methylated by ICMT (isoprenylcysteine carboxyl methyltransferase) (Simanshu *et al*., 2017). Some RAS isoforms, such as the human HRAS, NRAS and KRAS4A, carry a second type of lipidation — palmitoylation, which occurs on cysteine residues upstream of the CAAX motif (Hancock et al., 1989; Simanshu *et al*., 2017). HRAS and NRAS localize to Golgi membrane after prenylation followed by CAAX motif processing, then traffic to plasma membrane upon palmitoylation and go back to the Golgi membrane when the palmitoylation is erased by depalmitoylases (Campbell and Philips, 2021; Rocks et al., 2005). Membrane association mediated by lipidation is necessary for the activation and signaling of RAS. Both the palmitoylation and the depalmitoylation steps are potential targets for the treatment of cancers caused by HRAS or NRAS activating mutations, as evidenced by the findings that blocking either step disrupts the oncogenic activity of the NRAS G12D mutant (Remsberg et al., 2021; Zambetti et al., 2020).

In human, the palmitoylation of HRAS and NRAS is catalyzed by a protein complex formed by DHHC9 and GCP16 (also called GOLGA7) (Swarthout et al., 2005). DHHC9 belongs to a family of cysteine acyltransferases which catalyze the covalent attachment of a palmitoyl group (or other long-chain acyl groups) to the side chain of cysteine residues via thioester bond (Jiang et al., 2018). Members in this family share two structural features that are related to their conserved catalytic mechanism: one is a cysteine rich domain in which a conserved DHHC (Asp-His-His-Cys) motif has been identified as the catalytic center, thus the cysteine acyltransferases are also known as DHHC acyltransferases; the other is a long-chain acyl group-binding pocket formed by four transmembrane helices (Jiang *et al*., 2018). In human, 23 DHHC acyltransferases (DHHCs) have been identified to have various subcellular localization and substrate specificity (Jiang *et al*., 2018). The crystal structures of DHHC20 and its complex with palmitoyl-CoA have been reported, demonstrating the overall structure and the organization of the active site of DHHC20, which help understand the structures and catalytic activities of other DHHCs (Lee et al., 2022; Rana et al., 2018). In comparison with DHHC20 that functions as a monomer, DHHC9 is unique because it has to associate with GCP16 to acquire its catalytic activity and maintain its stability (Swarthout *et al*., 2005). But the molecular basis for the requirement of GCP16 by DHHC9 is not clear. It’s also not clear how the activity of the DHHC9-GCP16 complex is regulated.

The human DHHC9 and GCP16 were identified by searching the human genomic database with the protein sequences of yeast Erf2 and Erf4, respectively (Swarthout *et al*., 2005). The yeast Erf2-Erf4 complex is the first enzyme identified as a DHHC acyltransferase, which catalyzes the palmitoylation of RAS2 — a homolog of the human RAS proteins (Bartels et al., 1999; Lobo et al., 2002). Similar to the requirement of GCP16 by DHHC9, the accessory protein Erf4 is necessary for the catalytic activity and stability of Erf2 (Mitchell et al., 2012). It has been 20 years since the identification of the Erf2-Erf4 complex as a cysteine acyltransferase, however, its three-dimensional structure has not been reported.

In this study, we determined the cryo-EM structures of both the Erf2-Erf4 complex and the DHHC9-GCP16 complex, explaining why the accessory proteins Erf4 and GCP16 are required for the catalytic activity of Erf2 and DHHC9. In the cryo-EM structure, we identified a phospholipid binding to a positively charged patch formed by R85, R179 and R298 in DHHC9. The phospholipid functions as a hub to hold TM2, TM3 and a type II polyproline (PPII) helix of DHHC9 together. We also identified palmitoylation modifications at three cysteine residues, including C24, C25, and C288 in DHHC9, and showed that mutations of these residues significantly decreased the palmitoyltransferase activity of the DHHC9-GCP16 complex. In addition, we found that GCP16 also functioned as an accessory protein for DHHC14 and 18, and DHHC14 and 18 in complex with GCP16 could also catalyze the palmitoylation of HRAS and NRAS. These findings shed light on the molecular mechanism underlying RAS palmitoylation.

## Results

### Substrate specificity of RAS palmitoyltransferase

We overexpressed the Erf2-Erf4 complex and the DHHC9-GCP16 complex in insect cells and in HEK 293F cells, respectively, and purified them to homogeneity (Figure S1A and S1B). The ratio of zinc ions to DHHC9 was quantified using inductively coupled plasma mass spectrometry (ICP-MS), and the number was close to 2 (Figure S1C).

To evaluate their palmitoyltransferase activities, we used a fluorescent analog of palmitoyl-CoA (NBD-palmitoyl-CoA) as the palmitoyl group donor (Verardi et al., 2017), and used yeast RAS2 and human HRAS (and NRAS) as the substrates of Erf2 and DHHC9, respectively. Palmitoylation of RAS2 was observed in the presence of wild-type Erf2-Erf4 (lanes 9 and 10 in Figure 1A) but was not detectable either in the absence of Erf2-Erf4 (lanes 3 and 4 in Figure 1A) or in the presence of the C203A catalytic dead mutant of Erf2 in complex with Erf4 (lanes 7 and 8 in Figure 1A). C318 is the palmitoylation site in RAS2, downstream of which is the CAAX motif (Kuroda et al., 1993). Mutation of C318 to alanine decreased the palmitoylation level of RAS2, but a weak NBD signal was still observed, which may be due to the palmitoylation of C319 in the CAAX motif (lanes 5 and 6 in Figure 1A). Similarly, palmitoylation of HRAS and NRAS was observed in the presence of wild-type DHHC9-GCP16 but not the C169A catalytic dead mutant of DHHC9 in complex with GCP16 (lanes 7 to 10 in Figure 1B and Figure S1D). C181 and C184 in HRAS, and C181 in NRAS are the sites undergoing palmitoylation (Hancock *et al*., 1989); mutations of these cysteine residues to serine significantly decreased the palmitoylation level of HRAS and NRAS (lanes 5 and 6 in Figure 1B and Figure S1D). These results confirmed the enzymatic activities of the purified RAS palmitoyltransferases.

**Figure 1.**
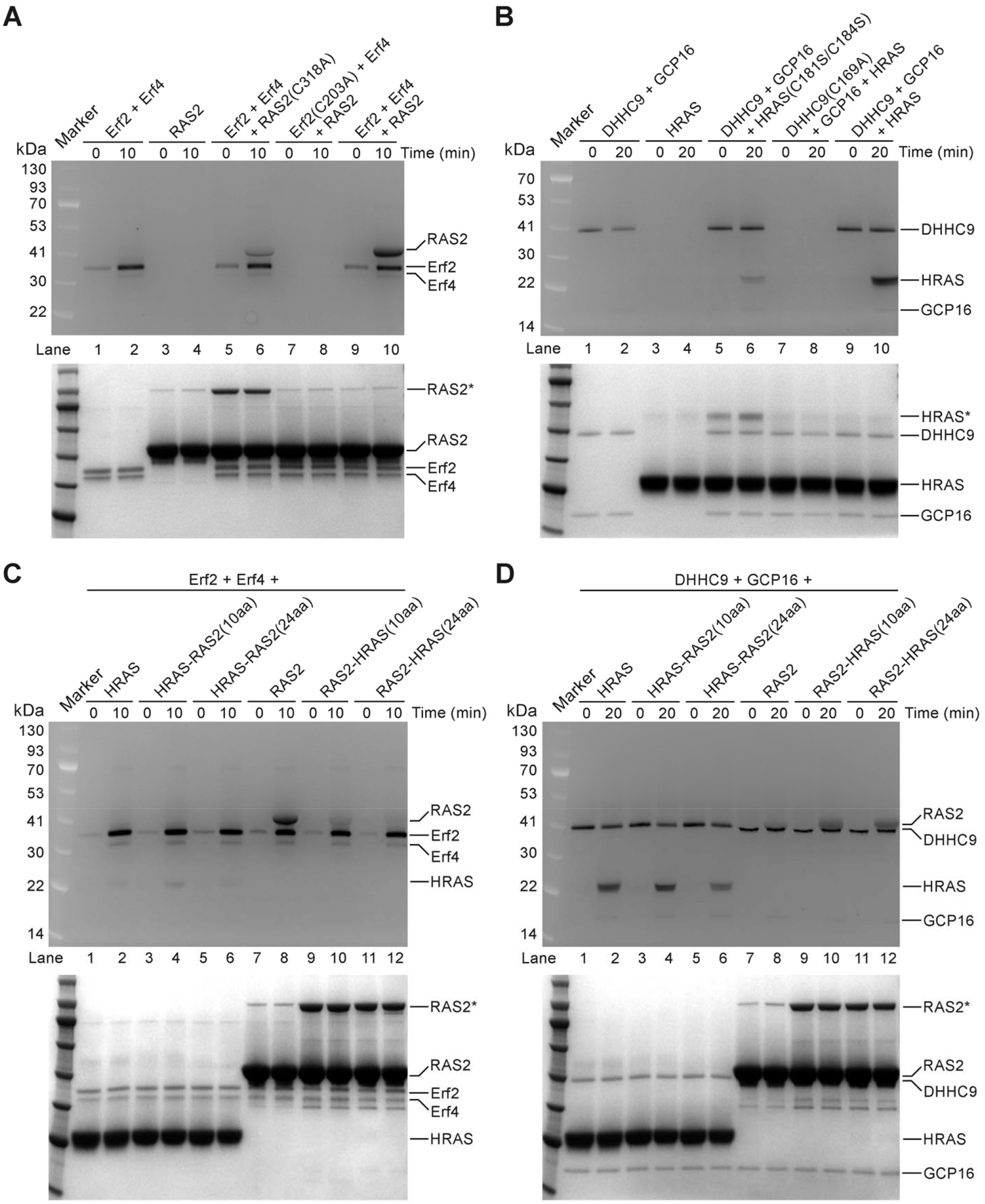
Substrate specificity of the RAS palmitoyltransferases. (A and B) The palmitoyltransferase activity of the Erf2-Erf4 complex (A) and that of the DHHC9-GCP16 complex (B) were evaluated using RAS2 and HRAS as the protein substrates, respectively, and using the NBD-palmitoyl-CoA as a substitute of palmitoyl-CoA. (C and D) The substrate specificity of the Erf2-Erf4 complex (C) and the DHHC9-GCP16 complex (D) were evaluated using chimeric RAS proteins. In each subfigure, the upper panel is the SDS-PAGE imaged by fluorescence imaging, representing the NBD-palmitoylation level, and the lower panel is the same SDS-PAGE visualized by Coomassie blue staining; the bands of RAS2 and HRAS indicated by asterisks were the dimers of RAS2 and HRAS, respectively.

Next, we tested whether the Erf2-Erf4 complex and the DHHC9-GCP16 complex could catalyze the palmitoylation of the substrates of each other. For HRAS, only a faint palmitoylation band was detected in the presence of the Erf2-Erf4 complex (lanes 1 and 2 in Figure 1C), which was much weaker than that in the presence of the DHHC9-GCP16 complex (lanes 1 and 2 in Figure 1D). Similar results were observed for RAS2, but in an opposite way — palmitoylation of RAS2 in the presence of the DHHC9-GCP16 complex was much weaker than that in the presence of the Erf2-Erf4 complex (lanes 7 and 8 in Figures 1C and 1D), consistent with the finding that expression of the human DHHC9-GCP16 complex in yeast only slightly reversed the growth defect caused by *erf2* gene knock-out (Mitchell et al., 2014).

The C-terminal region of HRAS was reported to be sufficient for the correct localization of HRAS in cells, and GFP with its C-terminus fused with the C-terminal 9, 19 or 24 amino acids (aa) of HRAS has been used to study the palmitoylation and trafficking of HRAS (Ahearn et al., 2011; Apolloni et al., 2000; Chai et al., 2013). To test the substrate specificity of the RAS palmitoyltransferases, we exchanged the C-terminal 10 or 24 aa between RAS2 and HRAS and checked the palmitoylation of the chimeric RAS proteins. When catalyzed by the Erf2-Erf4 complex, the palmitoylation of HRAS was at a very low level and had little increase when its C-terminal 10 or 24 aa was replaced by the corresponding sequence of RAS2 (lanes 3 to 6 in Figure 1C); the level of RAS2 palmitoylation was decreased a lot when its C-terminal 10 aa was replaced by the corresponding sequence in HRAS, and was further decreased when its C-terminal 24 aa was replaced (lanes 9 to 12 in Figure 1C). When catalyzed by the DHHC9-GCP16 complex, the level of HRAS palmitoylation was decreased but still detectable when its C-terminal 10 or 24 aa was replaced by the C-terminus of RAS2 (lanes 3 to 6 in Figure 1D); meanwhile, RAS2 palmitoylation increased to a level similar to that of HRAS when the C-terminal 10 or 24 aa of RAS2 was replaced by the corresponding sequence of HRAS (lanes 9 to 12 in Figure 1D). These data suggest that the C-terminus of RAS2 is necessary but not sufficient for its palmitoylation catalyzed by Erf2-Erf4; in contrast, the C-terminus of HRAS largely determines whether the whole protein can be palmitoylated or not with the catalysis of DHHC9-GCP16.

### Overall structures of RAS palmitoyltransferases

After validating the catalytic activity and substrate specificity of the purified Erf2-Erf4 complex and the DHHC9-GCP16 complex, we solved their structures using cryo-EM to a resolution of 3.5 Å and 3.4 Å, respectively (Figures S2 and S3, see also Table S1). In both structures, the DHHC enzyme (Erf2 or DHHC9) forms a heterodimer with the accessory protein (Erf4 or GCP16).

The overall structures of Erf2 and DHHC9 are similar to that of DHHC20 (Rana *et al*., 2018). Specifically, all of them have four transmembrane helices (TM), a large cytoplasmic region between TM2 and TM3 containing three anti-parallel β-sheets that form two zinc finger motifs (β3 to β8 in Erf2 and in DHHC9; β1 to β6 in DHHC20), and another cytoplasmic region after TM4 containing two conserved α-helices (α2 and α3 in Erf2 and in DHHC9; α3 and α4 in DHHC20) (Figures 2A, 2C, and 2E, and Figure S4A). The number of zinc ions in the DHHC9-GCP16 complex is consistent with the experimental data (Figure S1C).

**Figure 2.**
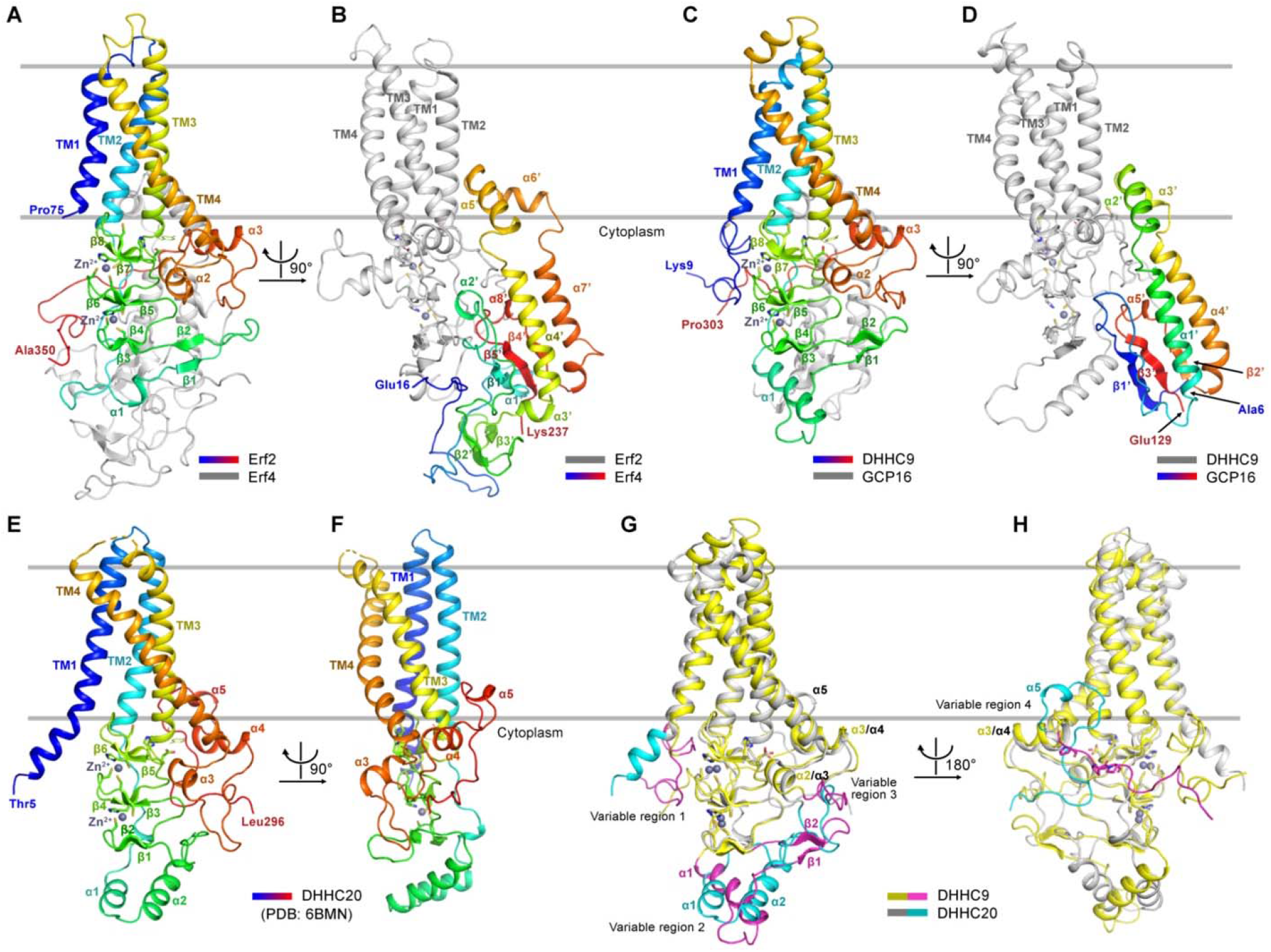
The overall structures of the RAS palmitoyltransferases. (A and B) The overall structure of the Erf2-Erf4 complex. The Erf2 in (A) and Erf4 in (B) are colored with a spectrum of colors from blue at the N-terminus to red at the C terminus. The Erf in (A) and Erf2 in (B) are colored gray. (C and D) The overall structure of the DHHC9-GCP16 complex. (E and F) The overall structure of DHHC20 (PDB code: 6BMN). (G and H) Alignment of the cryo-EM structure of DHHC9 in the DHHC9-GCP16 complex with the crystal structure of DHHC20 to identify regions that have distinct structures in DHHC9 and DHHC20 (defined here as variable regions). DHHC9 is colored gray, with the variable regions in it colored magentas; DHHC20 is colored yellow, with the variable regions in it colored cyan.

The C-terminus (residues 304-364) of DHHC9 is invisible in the cryo-EM structure, similarly to the C-terminus (residues 297-365) of DHHC20 in its crystal structure, suggesting their intrinsic flexibility (Rana *et al*., 2018). To understand the importance of the C-terminus in the palmitoyltransferase activity of DHHC9, we purified a truncated form (residues 1-305) of DHHC9 in complex with GCP16 and measured its activity. Our data showed that the palmitoyltransferase activity of the truncated DHHC9 was similar to that of the full-length DHHC9 (Figure S1E), suggesting that the C-terminal region may not contribute to the enzymatic activity of DHHC9.

The differences between the structures of DHHC9 and DHHC20 are mainly at four variable regions (Figures 2G and 2H, and Figure S4A). Firstly, the N-terminus (residues 9-32) of DHHC9 adopts a loop-like structure, in contrast, the N-terminus of DHHC20 is an α-helix which is an extension of TM1. Secondly, the region between TM2 and the zinc finger motifs in DHHC9 contains an α-helix (α1) and an anti-parallel β-sheet (β1 and β2), but in DHHC20 there are two α-helices (α1 and α2) (Figure 2A). Thirdly, the region between the two conserved α-helices after TM4 is a loop-like linker in both DHHC9 and DHHC20 but adopts distinct conformation. The fourth is the region after α3 in DHHC9 and that after α4 in DHHC20: in DHHC9 there is a type II polyproline (PPII) helix followed by a loop structure, but in DHHC20 there is a short α-helix followed by a loop structure with distinct orientation (Figure 2G and Figure S4A). The four variable regions in DHHC9, especially variable regions 1, 2 and 4, are important for the formation and function of the DHHC9-GCP16 complex, which will be discussed later.

In contrast to Erf2 and DHHC9, their accessory proteins Erf4 and GCP16 have no transmembrane helix, but both Erf4 and GCP16 have two α helices insert into the membrane (α5’ and α6’ in Erf4; α2’ and α3’ in GCP16) (Figures 2B and 2D). Interestingly, the two helices are at a position similar to that of helix α5 in DHHC20 (Figure 2F), which may partially explain why accessory proteins are required by Erf2 and DHHC9, but not DHHC20.

The structures of Erf4 and GCP16 also share other features. Both of them contain an anti-parallel β-sheets formed by three β-strands (β1’, β5’ and β4’ in Erf4; β1’, β3’ and β2’ in GCP16), and two α-helices in the cytoplasmic region (α4’ and α7’ in Erf4; α1’ and α4’ in GCP16) (Figures 2B and 2D, and Figure S4B). Erf4 has two unique segments in comparison with GCP16: one is a longer N-terminal region containing a long loop-like structure and a short helix (α1’), and the other is the long linker between β1’ strand and α4’ helix, which is folded into two β-strands (β2’ and β3’) (Figure 2B and Figure S4B).

### Structures of the active site of Erf2 and DHHC9

The core of the active site of DHHC acyltransferases is the conserved DHHC motif, in which the cysteine residue can accept an acyl group from acyl-CoA and then transfer it to cysteine residues in substrate proteins (Jiang *et al*., 2018; Rana *et al*., 2018). In the cryo-EM maps of Erf2 and DHHC9, we see density for aliphatic chain-like structures linked to the density of the catalytic cysteine in the DHHC motif, though we did not add acyl-CoA during protein purification and cryo-EM sample preparation. We tried to dock saturated acyl group with various chain-lengths into the density maps and found that only the saturated 14-carbon acyl group fit well (Figures 3A and 3B). The carbonyl oxygen of the acyl group forms hydrogen bonds (H-bonds) with the first histidine (H201 in Erf2; H167 in DHHC9) in the DHHC motif. Using an autoacylation assay, we found that both the Erf2-Erf4 complex and the DHHC9-GCP16 complex could use acyl CoA with chain lengths of 10, 14, 16 and 18-carbon as their substrates, but chain length longer than 18-carbon was not preferred (Figures 3C and 3D).

**Figure 3.**
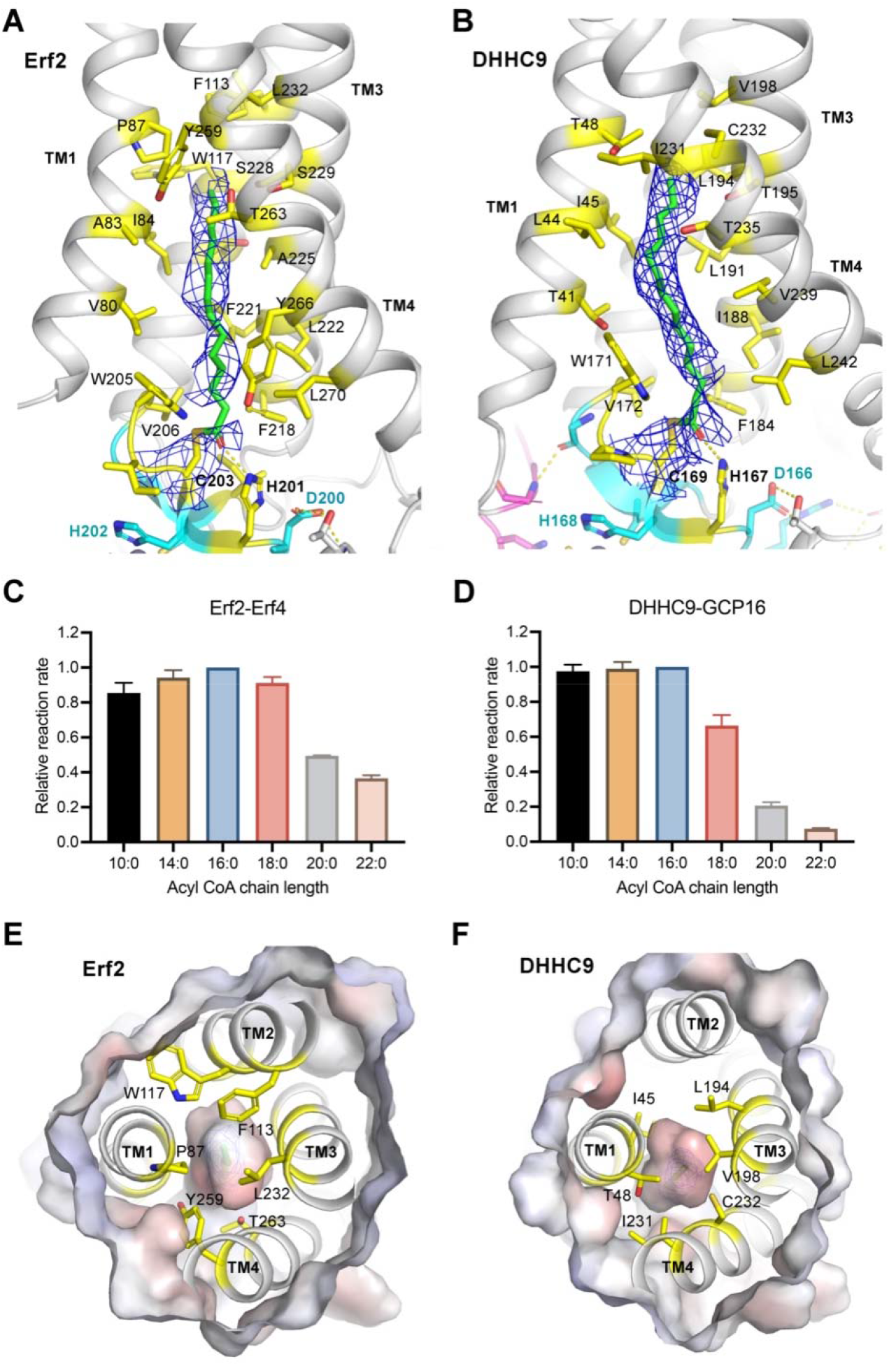
Organization of the active sites of the RAS palmitoyltransferases. (A and B) Structure of the DHHC motif and the acyl group-binding pocket of Erf2 (A) and that of DHHC9 (B). The acyl groups are showed as sticks and colored green. The cryo-EM map of the acyl group covalently linked to the catalytic cysteine is contoured at 4.0 σ (in Erf2) or 6.0 σ (in DHHC9) and colored blue. Residues within 6 Å of any atoms of the acyl group are showed as sticks and colored yellow. (C and D) The acyl CoA chain-length preference of the Erf2-Erf4 complex (C) and that of the DHHC9-GCP16 complex (D) measured by using an autoacylation assay. The data represent the mean ± SD of three independent measurements. (E and F) Top view of the acyl chain-binding pockets in Erf2 and DHHC9. The protein contact potential of Erf2 (E) and DHHC9 (F) were generated in PyMOL (version 2.4.2). The residues close to the tail of the acyl group are colored yellow.

The acyl group binding pockets in both Erf2 and DHHC9 consist mainly of hydrophobic residues from TM1, TM3 and TM4 (Figures 3A and 3B). In the pocket, residues close to the catalytic cysteine are highly conserved among Erf2, DHHC9 and DHHC20 (residues W205, V206, F218, F270 in Erf2 and the corresponding residues in DHHC9 and DHHC20), in contrast, residues around the tail of acyl chain are diverse (Figures 3E and 3F).

The DHHC motif is part of the cysteine-rich domain (CRD) that contains two zinc finger motifs (Figures 4A and 4D). The well-organized structure of the CRD enables the DHHC motif to adopt the right conformation to place the catalytic cysteine and the first histidine of the DHHC motif at the right position. Residues in the CRD are conserved among Erf2, DHHC9 and DHHC20 (Figure S4A). Each zinc finger motif has three cysteine residues and a histidine to coordinate the zinc ion. Disruption of the zinc finger motifs leads to a loss of function of the DHHC enzymes. For example, mutation R148W in DHHC9 identified in XLID patients was reported to affect the autoacylation of DHHC9 (Mitchell *et al*., 2014). We confirmed it in our assay (Figure S5A). Mutation of R148 to a bulky residue may disrupt the zinc-binding site formed by C141, C144, H154 and C161 (Figure 4D).

**Figure 4.**
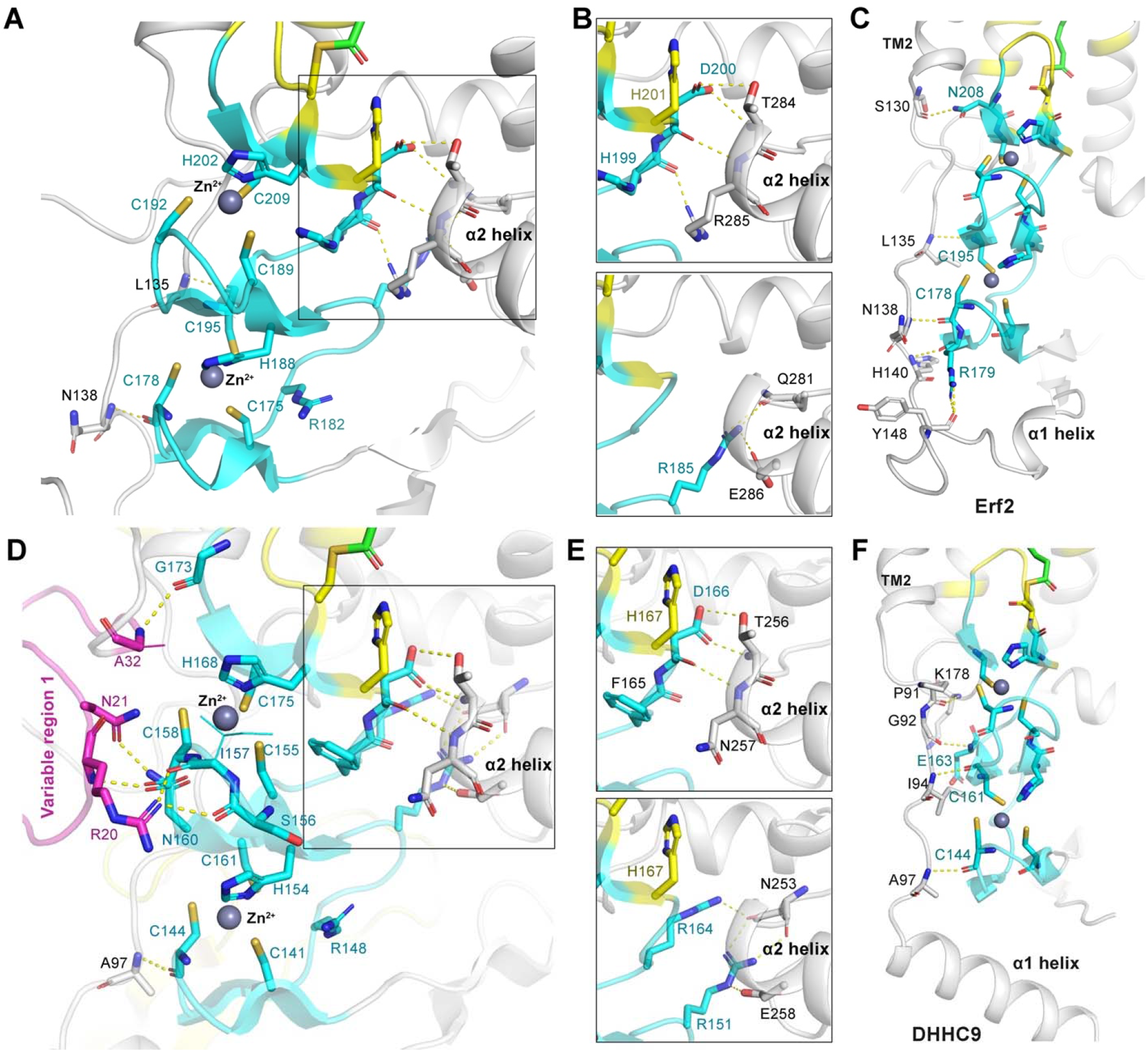
Organization of the zinc finger motifs of the RAS palmitoyltransferases. (A and D) Structures of the zinc finger motifs of Erf2 (A) and DHHC9 (D). The zinc finger motifs are colored cyan. The variable region 1 in DHHC9 is colored magenta. The residues that coordinate the zinc ions and that interact with the zinc finger motifs are showed as sticks. The hydrogen bonds are represented by yellow dash lines. (B and E) Interactions between the zinc finger motifs and the α2 helix in Erf2 (B) and that in DHHC9 (E). (C and F) Stabilization of the zinc finger motifs by the linker between TM2 and helix α1 in Erf2 (C) and that in DHHC9 (F).

The zinc finger motifs are stabilized by the surrounded structures. In both Erf2 and DHHC9, there are intense hydrogen bonding interactions between the zinc-binding motifs and the α2 helix (Figures 4A, 4B, 4D and 4E). In addition, the linkers between TM2 and helix α1 in both Erf2 and DHHC9 hold the zinc finger motifs together through hydrogen bonding (Figures 4C and 4F). In DHHC9, the variable region 1 is also involved in stabilizing the zinc finger motifs: residues R20, N21 and A32 in the variable region 1 form five H-bonds with the main chains of S156, I157, C158, G173 and the side chain of N160 (Figure 4D). Mutation of R20 to glutamate decreased the activity of DHHC9 (Figure S5B), supporting the importance of the variable region 1.

### Interactions between DHHC9 and its accessory protein GCP16

DHHC9 interacts with GCP16 mainly through four interfaces (Figure 5A). One is between TM2, TM3 helices of DHHC9 and α3’ helix of GCP16 (Figure 5B). The side chain of R85 in DHHC9 TM2 donates a H-bond to the main chain of Y76 in GCP16 α3’ helix; Y183 in DHHC9 TM3 interacts with Y76 in GCP16 α3’ helix through π–π stacking. Mutation of either DHHC9 R85 or GCP16 Y76 to alanine decreased the catalytic activity of the DHHC9-GCP16 complex (Figure S5C). The second is between the PPII helix close to the C-terminus of DHHC9 and the α-helices of GCP16 (Figure 5C). P290 and P293 in the DHHC9 PPII helix dock into two weakly negatively charged pockets formed by α1’-α2’ and α4’-α5’ helices of GCP16, while P292 in the PPII helix form a CH-π hydrogen bond with Y86 in GCP16 α3’ helix (Nishio et al., 2014). The first and second interfaces stabilize DHHC9 by holding its TM2, TM3 and the PPII helix together.

**Figure 5.**
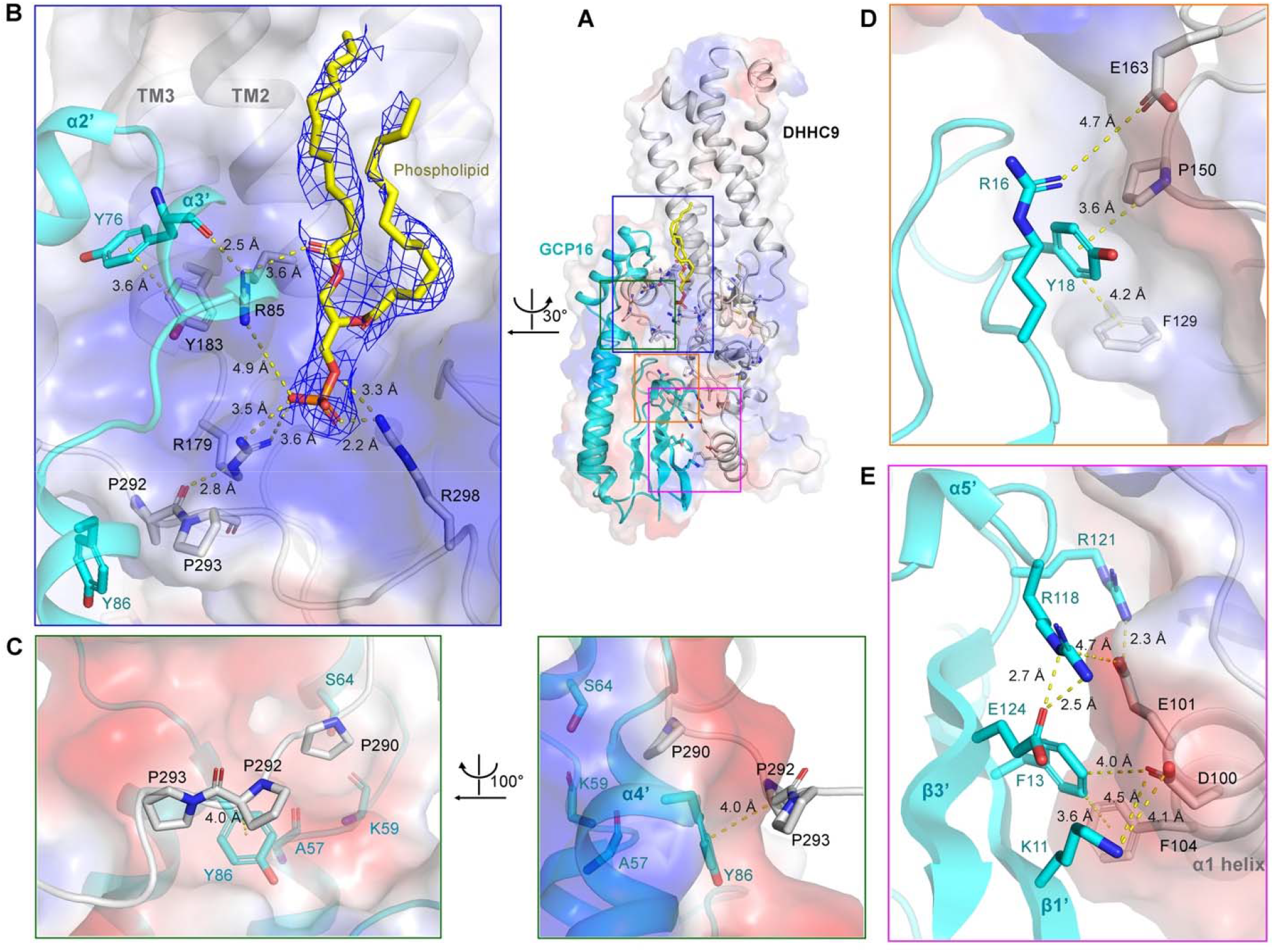
Interactions between DHHC9 and GCP16. (A) The overall structure of the DHHC9-GCP16 complex. (B) R85 and Y183 in TM2 and TM3 of DHHC9 interact with Y76 in the α3’ helix of GCP16. A phospholipid was identified to interact with three arginine residues in DHHC9 (R85, R179 and R298). The cryo-EM map of the phospholipid is contoured at 6.0 σ and colored blue. The hydrogen bonds are represented by yellow dash lines. (C) Three proline residues (P290, P292 and P293) in the PPII helix of DHHC9 dock into a groove in GCP16. (D) P150 and E163 in the zinc finger motifs and F129 of DHHC9 interact with R16 and Y18 in the loop following GCP16 β1’ strand. (E) Residues in the α1 helix of DHHC9 interact with residues in the β1’, β3’ strands and in a loop following the α5’ helix of GCP16.

The third is between the zinc finger motifs of DHHC9 and the loop following the β1’ strand of GCP16 (Figure 5D). P150 and F129 in DHHC9 interact with Y18 in GCP16 through CH-π interaction and through π–π stacking, respectively; E163 in DHHC9 interacts with R16 in GCP16 through charge−charge interaction. The importance of DHHC9 P150 is consistent with the identification of P150S mutation in DHHC9 as a cause of X-linked intellectual disorders (XLIDs) (Mitchell *et al*., 2014). We also found that the DHHC9 P150S mutant had a decreased autopalmitoylation level and lost its catalytic activity (Figure S5A). The fourth interface is between the α1 helix of DHHC9 and residues from the β1’, β3’ strands and a loop following the α5’ helix of GCP16 (Figure 5E). DHHC9 D100 interacts with K11 and F13 in GCP16 through charge-charge interaction and anion-π interaction, respectively; DHHC9 E101 interacts with R118 and R121 in GCP16 through charge-charge interaction and charge-stabilized H-bond, respectively; in addition, DHHC9 F104 forms π–π stacking with GCP16 F13. The fourth interface stabilizes DHHC9 α1 helix, thus indirectly stabilize the linker between TM2 and the α1 helix which interacts with the zinc finger motifs through H-bonds (Figure 4F). Taken together, the third and fourth interfaces stabilize the zinc finger motifs of DHHC9.

An important finding in the DHHC9-GCP16 structure is the presence of a phospholipid around a positively charged patch formed by R85, R179 and R298 in DHHC9 (Figure 5B). Based on the density of the cryo-EM map, we docked a 1,2-Dilauroyl-sn-glycero-3-phosphate (DLPA) into the structure. The phosphate group of the phospholipid interacts with R85, R179 and R298 through charge-charge interactions and H-bonds; the carbonyl oxygen of one of the two lauroyl groups of the phospholipid interacts with R85 through charge-charge interaction; R179 donates a hydrogen bond to the main chain of P292 in the DHHC9 PPII helix. Either the R85A or the R298E mutation decreased the palmitoyltransferase activity of DHHC9, and the R298E mutation also decreased the autopalmitoylation level of DHHC9 (Figures S5C and S5D). These data suggest phospholipid functions as a hub to stabilize the conformation of TM2, TM3 and the PPII helix of DHHC9.

The interactions between Erf2 and Erf4 share similar pattern with that between DHHC9 and GCP16 but are different in details (Figure S6). Firstly, R126 and R216 in TM2 and TM3 of Erf2 interact with G176, S179 and N164 in α5’ and α6’ helices of Erf4 through H-bonds; P325 and P328 in a PPII helix similar to that in DHHC9 dock into a groove formed between α5’-α6’ and α4’-α7’ helices (Figure S6B). Secondly, T162, H163, S165 and I166 in a loop between β1 and β2 strands of Erf2 accept four H-bonds from R17 and R83 of Erf4 (Figure S6C). Thirdly, residues between the TM2 and β1 strand of Erf2 (L141, Q143, L144, R145, P151, E153 and Y154) interact with residues in the N-terminal region (Y23, H24, F26 and S27) and the loop between the β2’ and β3’ strands (Y117 and D118) of Erf4 through hydrogen bonding, π–π stacking and CH-π interactions (Figures S6D and S6E), facilitating the binding of the linker between TM2 and α1 helix of Erf2 with the zinc finger motifs (Figure 4C).

### Palmitoylation regulates DHHC9 activity

In the cryo-EM map of the DHHC9-GCP16 complex, there are extra densities linked to the side chains of C24, C25 and C288 in DHHC9, and the shape of the densities indicates that these cysteine residues were modified by acyl groups (Figures 6A and 6B). Using mass spectrometry (MS), we identified palmitoylation of DHHC9 at both C25 and C288 (Figures S7A and S7B); though no palmitoylation at C24 was identified by MS, a palmitoyl group covalently linked to C24 matches well with the cryo-EM density (Figure 6A).

**Figure 6.**
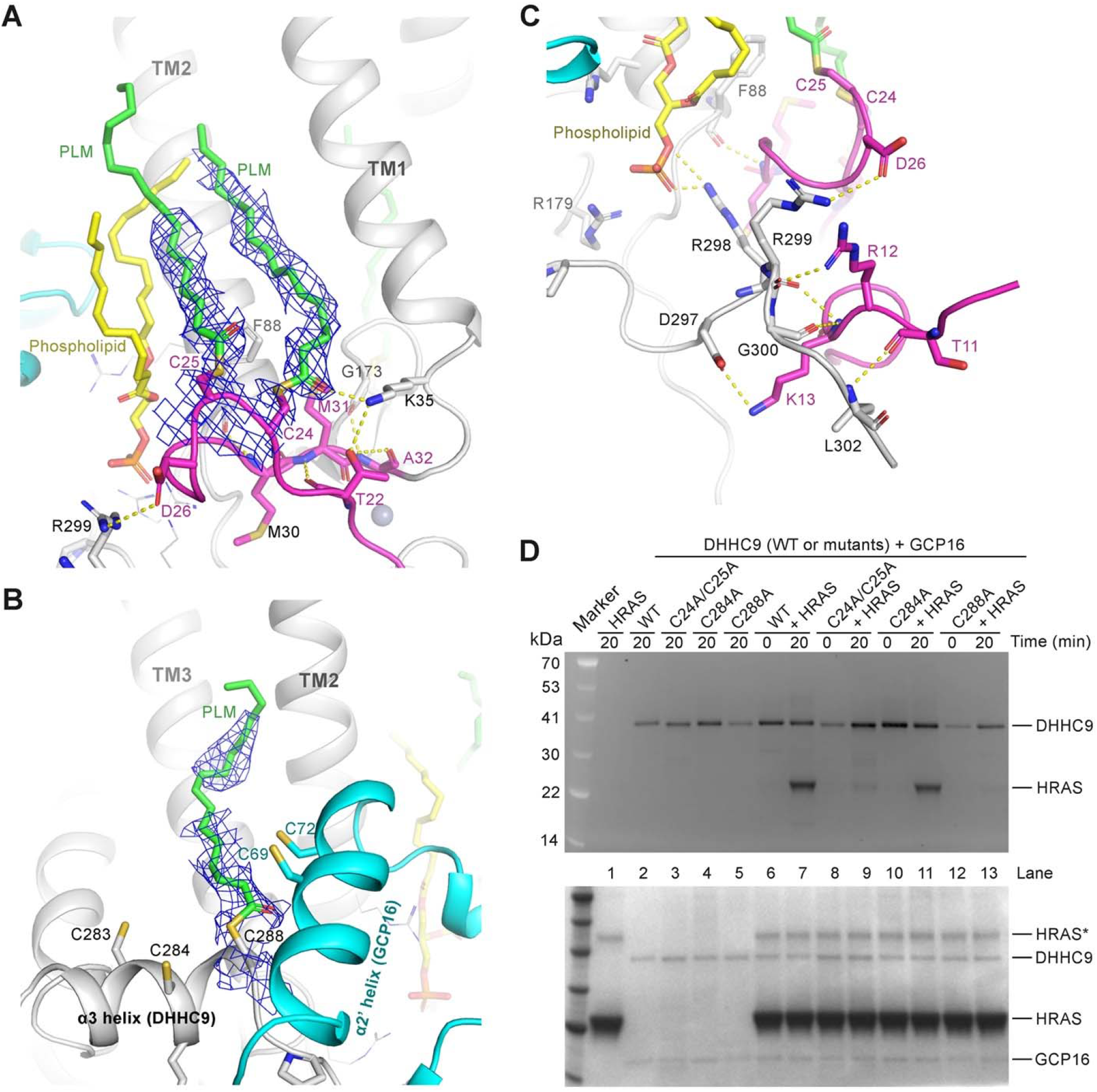
Cysteine palmitoylation regulates DHHC9 catalytic activity. (A and B) Identification of palmitoylation sites in DHHC9. Extra density was identified along the side chains of C24, C25, and C288. The cryo-EM map of the palmitoyl groups is contoured at 6.0 σ and colored blue. The palmitoyl groups are showed as sticks and colored green. The hydrogen bonds are represented by yellow dash lines. (C) Interactions between the region 1 and residues following the PPII helix of DHHC9. (D) Evaluation of the effect of mutating C24, C25, and C288 in DHHC9 to alanine on the acyltransferase activity of the DHHC9-GCP16 complex. The upper panel is the SDS-PAGE imaged by fluorescence imaging, representing the NBD-palmitoylation level, and the lower panel is the same SDS-PAGE visualized by Coomassie blue staining; the bands of HRAS indicated by an asterisk were the dimer of HRAS.

C24 and C25 are in a loop of the variable region 1 in DHHC9 and is close to the phospholipid (Figure 6A). The variable region 1 interacts with multiple regions in DHHC9. Firstly, the side chain of T22 and the carbonyl oxygen of the palmitoyl group attached to C24 form H-bonds with the side chain of K35 at the N-terminus of TM1; the main chain of M30 accepts a H-bonds from the main chain of F88 in TM2. Secondly, the variable region 1 also interacts with the zinc finger motifs (Figure 4D). Thirdly, T11, R12, K13 and D26 in the variable region 1 form H-bonds with residues D297, R298, R299, G300 and L302, thus keeping the PPII helix of DHHC9 at a proper position to facilitate its interaction with GCP16 (Figure 6C). Palmitoylation of C24 and C25 anchors the variable region 1 to the lipid bilayer, stabilizing the conformation of the variable region 1 and enabling it to interact with the multiple regions in DHHC9. Mutation of C24 and C25 to alanine showed little effect on the autopalmitoylation level of DHHC9 but severely decreased the palmitoyltransferase activity of the DHHC9-GCP16 complex (Figure 6D).

C288 at the C-terminus of DHHC9 α3 helix is surrounded by TM2 and TM3 of DHHC9 and helix α2’ of GCP16 (Figure 6B). The palmitoyl group attached to C288 inserts into the lipid bilayer and docks into a hydrophobic pocket between TM2 and TM3 to bring DHHC9 α3 helix close to TM2 and TM3. The C288A mutant lost the activity to catalyze HRAS palmitoylation (Figure 6D). C288 is at the center of a cysteine cluster including C283, C284 and C288 in DHHC9 α3 helix and C69 and C72 in GCP16 α2’ helix. The activity of the DHHC9 C284A mutant was similar to that of wild-type DHHC9 (Figure 6D), but the C69A/C72A double mutant of GCP16 in complex with DHHC9 showed a decreased acyltransferase activity (Figure 6D and Figure S5C). Though we did not see palmitoylation at C69 or C72 of GCP16 in our MS data, a previous study showed that the two cysteine residues were palmitoylated, and the C69A/C72A double mutant could not localize to the Golgi (Ohta et al., 2003). These results suggest that membrane attachment of DHHC9 α3 helix and GCP16 α2’ helix through cysteine palmitoylation is important for maintaining the stability of the DHHC9-GCP16 complex.

### DHHC14 and 18 also form complexes with GCP16 to catalyze RAS palmitoylation

Alignment of the protein sequence of DHHC9 with that of the other 22 human DHHCs using the BLAST (Basic Local Alignment Search Tool) (Johnson et al., 2008) shows that DHHC14, 18, 5, and 8 are most closely related to DHHC9 (Figure S7C), and DHHC19 also has a high alignment score. A PPII helix in DHHC9 locates at the center of the DHHC9-GCP16 interface. We speculate that this PPII helix is a unique feature of DHHCs which require accessory proteins similar to GCP16. Two important proline residues (P290 and P293) in this PPII helix are conserved in Erf2 but not in DHHC20, and also conserved in DHHC14, 18, 8 and 19 (Figure S7D); the residue in DHHC5 corresponding to DHHC9 P290 is a S247, but after S247 there is a proline residue (Figure S7D). The alignment suggests that DHHC14, 18, 8, 5 and 19 may also have GCP16-like accessory proteins. Indeed, GCP16 has been reported to also form complexes with DHHC5 and DHHC8 (Ko et al., 2019); a homolog of GCP16 — GOLGA7B also interacts with DHHC5 to regulate its function (Woodley and Collins, 2019).

To test whether DHHC14 and 18 can form complex with GCP16, we transiently expressed them alone or together with GCP16 in Expi293F cells. In consistent with our speculation, DHHC14 and 18 could be co-purified with GCP16 (Figures 7A and 7B), but formed aggregates when each of them was expressed alone (data not shown). We also co-expressed DHHC19 with GCP16 in Expi293F cells, but unfortunately no target protein was obtained after purification.

**Figure 7.**
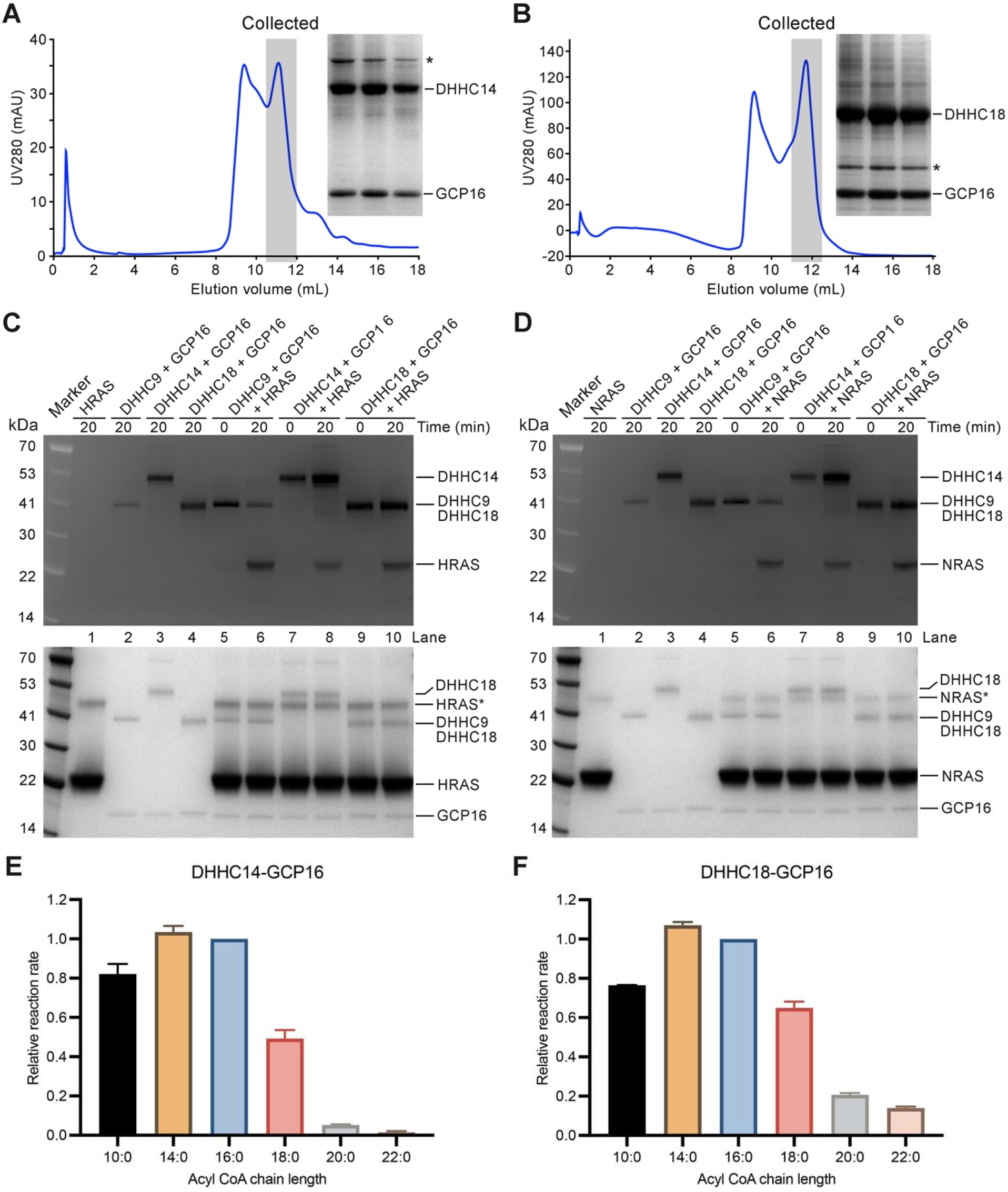
GCP16 is an accessory protein of DHHC14 and 18. (A and B) Size exclusion chromatography (SEC) of GCP16 in complexes with DHHC14 (A) and DHHC18 (B). GCP16 with a N-terminal His tag and DHHC14 or 18 with a C-terminal FLAG tag were transiently co-expressed in Expi293F cells and tandemly purified by anti-His tag and anti-FLAG tag affinity resins and then further purified by a SEC column (Superdex 200 Increase). The peak fractions were collected and analyzed by SDS-PAGE gels. The gels were stained by Coomassie Blue. The band indicated by asterisk in (A) was supposed to be a chaperone. The band indicated by asterisk in (B) is being identified. (C and D) Palmitoylation of HRAS (C) and NRAS (D) catalyzed by DHHC9, 14 and 18 in complex with GCP16. The upper panel is the SDS-PAGE imaged by fluorescence imaging, representing the NBD-palmitoylation level, and the lower panel is the same SDS-PAGE visualized by Coomassie blue staining; the bands of HRAS or NRAS indicated by an asterisk were the dimer of HRAS or NRAS. (E and F) The acyl CoA chain-length preference of DHHC14 and 18 in complex with GCP16. The data represent the mean ± SD of three independent measurements.

Next, we measured the palmitoyltransferase activity of DHHC14 and 18 in complex with GCP16. Both of them were able to catalyze the palmitoylation of HRAS and NRAS (Figures 7C and 7D), and prefer acyl CoA with chain lengths of 14 and 16-carbon (Figure 7E and 7F). These data suggest that GCP16 is a universal accessory protein for DHHC9, 14 and 18, and DHHC14 and 18 are also involved in the regulation of RAS signaling.

## Discussion

Catalytic mechanism, substrate specificity, and regulation are the focus of enzyme studies. For the DHHC acyltransferases, a two-step ping-pong catalytic mechanism has been proposed: the acyl group is first transferred from acyl-CoA to the cysteine in the DHHC motif and then transferred to cysteine residues in substrate proteins (Jennings and Linder, 2012; Mitchell et al., 2010). The structures of DHHC20 in the lipid-free state, the covalent inhibitor 2-bromopalmitate (2-BP)-bound state, and the palmitoyl-CoA-bound state have been solved (Lee *et al*., 2022; Rana *et al*., 2018). The Erf2-Erf4 and the DHHC9-GCP16 complexes in our structures have their catalytic cysteines modified by a 14-carbon acyl group, therefore, our structures represent the first autoacylated structures of DHHCs and indicate that the RAS palmitoyltransferases may prefer acyl group with a chain-length of 14 carbon (Figures 3A and 3B). As for the specificity of the RAS palmitoyltransferases for protein substrates, the Erf2-Erf4 complex showed very weak activity to catalyze acylation of the substrate of the DHHC9-GCP16 complex, and vice versa (Figures 1C and 1D). This finding is different from the conclusion from a previous study which claims that no specific protein acyltransferase activity exists for RAS (Rocks et al., 2010). The Erf2-Erf4 complex could not catalyze the acylation of HRAS-RAS2(24aa), but the DHHC9-GCP16 complex could catalyze the acylation of RAS2-HRAS(24aa) (Figures 1C and 1D), indicating that the substrate specificity of DHHC9-GCP16 is mainly determined by its interaction the C-terminal 24 aa of HRAS, but for Erf2-Erf4, regions beyond the C-terminal 24 aa of RAS2 also contribute to the specific interaction between Erf2-Erf4 and RAS2.

The importance of acylation in RAS signaling has been known for more than 30 years (Hancock *et al*., 1989), however, the mechanism for the regulation of RAS acylation is poorly understood. Among the 23 human DHHCs, most of them function as a monomer, but a few have accessory proteins (Salaun et al., 2020). The requirement of GCP16 by DHHC9 has been demonstrated by previous studies (Mitchell *et al*., 2014; Swarthout *et al*., 2005). Our structures show that GCP16 (or Erf4) binds to a surface of DHHC9 (or Erf2) that is opposite to the DHHC motif (Figures 2A to 2D), suggesting that GCP16 and Erf4 are not directly involved in the catalytic process, but stabilize the architecture of DHHC9 and Erf2, respectively. Furthermore, our study and previous studies (Ko *et al*., 2019) demonstrate that GCP16 is an accessory protein for at least 5 of the 23 human DHHC acyltransferases (DHHC5, 8, 9, 14, and 18), indicating that GCP16 plays a pivotal role in the regulation of DHHC acyltransferases.

In the cryo-EM structure of the DHHC9-GCP16 complex, we identified the binding of a phospholipid to three arginine residues in DHHC9 (Figure 5A). Among the three arginine residues, R85 is only conserved in DHHC14; R179 is conserved in DHHC14, DHHC18, DHHC8, DHHC5, and DHHC19; R298 is conserved in DHHC14, DHHC18, and DHHC5 (Figure S7D). Therefore, we speculate that at least DHHC14 also binds to a phospholipid similar to that in DHHC9.

Another interesting finding is the palmitoylation of three cysteine residues in DHHC9. Among the three cysteine residues, C25 is conserved in DHHC14 and 18, while C288 is conserved in DHHC14, 18, 8, 5, and 19, suggesting a conserved role of cysteine palmitoylation in the regulation of DHHCs. In consistent with our finding, the corresponding residue of C288 in DHHC5, C245, was reported to be palmitoylated together with C236 and C237 in DHHC5 (Yang et al., 2010), and mutating the three cysteine residues to serine or alanine disrupted the interaction between DHHC5 and GCP16 or GOLGA7B (Ko *et al*., 2019; Woodley and Collins, 2019). It is not clear the palmitoylation of C24, C25 and C288 in DHHC9 is a non-enzymatic event or catalyzed by DHHC9 or other DHHCs. The only palmitoylation cascade reported so far is the palmitoylation of DHHC6 by DHHC16 (Abrami et al., 2017). Identification of the DHHCs that catalyze the palmitoylation of DHHC9 would reveal more palmitoylation cascades.

### Limitations of the study

First, we showed that the Erf2-Erf4 complex and the DHHC9-GCP16 complex had different protein substrate selectivity, but did not provide an explanation. Three-dimensional structures of DHHC-substrate complexes are required to elucidate how RAS palmitoyltransferases recognize their substrates. Second, the phospholipid in DHHC9 was treated as DLPA in the cryo-EM structure, but whether other phospholipids can also bind to DHHC9 and whether the phospholipid is a constitutive component of the DHHC9-GCP16 complex or plays a regulatory role are not clear. Third, we showed that GCP16 also formed complexes with DHHC14 and 18 to catalyze the palmitoylation of HRAS and NRAS using *in vitro* assays. The roles of DHHC14 and 18 in the RAS signaling need to be further studied using cell-based and *in vivo* assays.

## Acknowledgments

We thank Dr. Peilong Lu and Ke Sun for their help with structure prediction; the Cryo-EM Facility and HPC Center of Westlake University for providing cryo-EM and computation support; the Mass Spectrometry & Metabolomics Core Facility at the Center for Biomedical Research Core Facilities of Westlake University for sample analysis; Yinjuan Chen and Ke Wang at the Instrumentation and Service Center for Molecular Sciences at Westlake University for the ICP-MS analysis. This work was supported by Central Guidance on Local Science and Technology Development Fund (2022ZY1006) and Westlake Education Foundation (Q.H., J.W.).

## Data and materials availability

The cryo-EM structures have been deposited in the Protein Data Bank (www.rcsb.org) with the accession codes 8HFC (Erf2-Er4 complex) and 8HF3 (DHHC9-GCP16 complex). The cryo-EM maps have been deposited in the Electron Microscopy Data Bank with the accession codes EMD-34717 (Erf2-Er4 complex) and EMD-34711 (DHHC9-GCP16 complex). All other data are available in the manuscript or the supplementary materials.

## Author contributions

Q.H. conceived and supervised the project; A.Y. and S.L. purified the proteins, performed the biochemical assays, prepared the cryo-EM samples, and collected the cryo-EM data; Y.Z. and J.W. calculated the cryo-EM map and built the model; J.C. and S.F. performed the mass spectrum experiment; all authors contributed to data analysis; Q.H. and J.W. wrote the manuscript.

