## Supplementary Information for "Regulation of RAS palmitoyltransferases by accessory proteins and palmitoylation"

**Method details**

Genes and cloning

The genes of yeast palmitoyltransferase Erf2 (GenBank: NM_001182133.1) and its accessory protein Erf4 (GenBank: NM_001183364.1) were isolated from *Saccharomyces cerevisiae* S288C using Q5 High-Fidelity DNA Polymerase (NEB) and cloned into a modified pFastBac baculovirus expression vector. The construct of Erf2 containes an N-terminal 12x His tag followed by a DrICE cleavage site (AspGluValAsp↓Ala). Site-directed mutagenesis was performed to mutate Cys203 to alanine to get the Erf2(C203A) expression construct. The construct of Erf4 contained an N-terminal 2x FLAG tag followed by a DrICE cleavage site. The construct harboring Erf2 or the C203A mutant was co-expressed with Erf4 in *Spodoptera frugiperda* (Sf9) cells. Cell cultures were grown in Sf-900™ II SFM medium (Gibco) to a density of 1.5 - 2 million cells per mL and then infected with two separate baculoviruses at a volume ratio of 1:1 for Erf2 and Erf4. After infection, the cells were cultured at 27 °C for 48 h, and then collected by centrifugation and stored at -80°C until use.

The CDS regions of the genes of human DHHC9 (GenBank: NM_016032.4) and DHHC18 (GenBank: NM_032283.3) were chemically synthesized (Genewiz); the cDNA of human DHHC19 (GenBank: NM_001039617.2) was purchased from YouBio; the gene of human DHHC14 (GenPept: NP_078906.2) was codon-optimized and chemically synthesized (Genewiz). The DHHC genes were cloned into pCAGGS vector with a C-terminal FLAG tag. The cDNA of human GCP16 (GenBank: NM_001002296.2) was purchased from YouBio and its CDS region was cloned into the pcDNA3.1 vector with an N-terminal 6x His tag. Site directed mutagenesis was performed to generate the expression constructs of the mutants of DHHC9 or GCP16. The plasmid harboring DHHC9, 14, 18, or 19 was co-transfected with the plasmid of GCP16 in Expi293F cells (Invitrogen). Cell cultures were grown in SMM 293-TI Expression Medium (Sino Biological) at 37 °C with 5% CO_2_ to a density of 2 million cells per mL. For each liter cell culture, 1 mg of DHHC plasmid and 1 mg of GCP16 plasmid were pre-mixed at a molar ratio of 1:1 and transfected with polyethylenimines (PEIs, Polysciences). After transfection, the cells were cultured at 37 °C for 48 h, and then collected by centrifugation and stored at -80 °C until use.

The yeast *RAS2* (GenBank: NM_001182936.1) gene, the CDS region of human *HRAS* (GenBank: NM_005343.4) or the *NRAS* (GenBank: NM_002524.5) gene was cloned into the pET15b vector with a N-terminal 6x His tag followed by a thrombin cleavage site (RAS2) or a DrICE cleavage site (HRAS and NRAS). Site directed mutagenesis was performed to generate the expression constructs of the mutants of RAS2, HRAS and NRAS. The C-terminal amino acid exchanged plasmids were constructed using KOD OneTM PCR Master Mix (TOYOBO). Plasmids were transformed into BL21(DE3) and the cells were cultured at 37 °C until OD_600_ reached 0.8-1.0, then the temperature was lowered to 20 °C for 1 hour and the expression was induced by isopropyl β-D-1-thiogalactopyranoside (IPTG) with a final concentration of 0.5 mM for 12-18 h. The cells were then collected by centrifugation and stored at -80°C until use.

Protein expression and purification

The insect cells containing the transiently expressed Erf2-Erf4 complex were resuspended in lysis buffer (25 mM Tris-HCl pH 8.0, 150 mM NaCl) supplemented with a protease inhibitor cocktail (MCE) and PMSF, and then incubated with 2% (w/v) n-Dodecyl-β-D-Maltopyranoside (DDM, Anatrace) for 3 h at 4 °C. The lysed cells were centrifuged at 100,000 g for 1 h at 4 °C, then the supernatant was collected and incubated with TALON resin (TaKaRa) for 2 h at 4 °C. The resin was washed with 15 column volumes of wash buffer (25 mM Tris-HCl pH 8.0, 150 mM NaCl and 0.02% DDM) followed by 5 column volumes of wash buffer supplemented with 25 mM imidazole. The Erf2-Erf4 complex was eluted with 5-6 column volumes of elution buffer (20 mM Tris-HCl pH 8.0, 150 mM NaCl, 0.02% DDM and 300 mM imidazole), and incubated with anti-FLAG tag affinity resin (Genscript) for 2 h at 4 °C. After washing the resin with 15 column volumes of wash buffer (25 mM Tris-HCl pH 8.0, 150 mM NaCl and 0.02% DDM), the Erf2-Erf4 complex was eluted with 6 column volumes of elution buffer (20 mM Tris-HCl pH 8.0, 150 mM NaCl, 0.02% DDM and 250 µg/ml FLAG peptide). The eluate was concentrated with a 50 kDa molecular weight cut-off centrifugal filter unit (Millipore Corporation), and further purified by a Superdex 200 Increase 10/300 GL (Cytiva) column with the buffer containing 25 mM Tris-HCl, pH 8.0, 150 mM NaCl, 0.02% DDM. The peak fractions were collected and used for the acyltransferase assays and cryo-EM studies.

The Expi293F cells containing the transiently expressed DHHC-GCP16 complex were resuspended in lysis buffer (50 mM Tris-HCl pH 7.5, 150 mM NaCl, 10% glycerol, and 5 mM β-ME) supplemented with a protease inhibitor cocktail (MCE) and PMSF, then lysed by a microfluidizer. The cell lysate was centrifuged at 150,000 g for 1 hour at 4 °C. The cell pellet was resuspended in lysis buffer containing 1% DDM and 0.2% Cholesteryl Hemisuccinate Tris Salt (CHS, Anatrace), incubated overnight at 4 °C, then centrifuged at 150,000 g for 1 h at 4 °C. The supernatant was incubated with Ni-NTA resin (Thermo Fisher) for 2 h at 4 °C. Then the resin was washed with 15 column volumes of wash buffer (50 mM Tris-HCl pH 7.5, 150 mM NaCl, 10% glycerol, 5 mM β-ME, 0.02% DDM and 0.004% CHS), and 5 column volumes of the wash buffer supplemented with 5 mM imidazole. The DHHC-GCP16 complex was eluted with 6-8 column volumes of elution buffer (50 mM Tris-HCl pH 7.5, 150 mM NaCl, 10% glycerol, 5 mM β-ME, 0.02% DDM, 0.004% CHS and 300 mM imidazole). The eluate was incubated with anti-FLAG tag affinity resin for 2 h at 4 °C, and then the resin was washed with 15-20 column volumes of the wash buffer (50 mM Tris-HCl pH 7.5, 150 mM NaCl, 10% glycerol, 5 mM β-ME, 0.02% DDM and 0.004% CHS). The protein was eluted with 6-8 column volumes of elution buffer (50 mM Tris-HCl pH 7.5, 150 mM NaCl, 10% glycerol, 5 mM β-ME, 0.02% DDM, 0.004% CHS and 250 µg/ml FLAG peptide). The eluate was concentrated with a 10 kDa molecular weight cut-off centrifugal filter unit and further purified by a Superdex 200 Increase 10/300 GL column with the buffer containing 50 mM Tris-HCl, pH 7.5, 125 mM NaCl, 0.5 mM dithiothreitol (DTT), 0.02% DDM and 0.004% CHS. The peak fractions were collected and used for substrate acyltransferase assays and cryo-EM studies.

For the purification of RAS2, the BL21(DE3) cells expressing RAS2 were resuspended in lysis buffer (25 mM Tris-HCl pH 8.0, 150 mM NaCl) supplemented with PMSF, and disrupted by sonication on ice. The lysed cells were centrifuged with a speed of 12,000 rpm at 4 °C for 1 h, then the supernatant was incubated with TALON resin for 2 h at 4 °C. The resin was washed with 15 column volumes of the lysis buffer, followed by 5 column volumes of wash buffer (lysis buffer plus 10 mM imidazole). The protein was eluted with 8-10 column volumes of elution buffer (25 mM Tris-HCl pH 8.0, 150 mM NaCl and 250 mM imidazole). The eluate was diluted using buffer A (25 mM Tris-HCl pH 8.0, 1 mM MgCl_2_), then loaded onto a SOURCE 15Q (Cytiva) column and eluted by a linear gradient from 100% buffer A to 60% buffer B (25 mM Tris-HCl pH 8.0, 1 mM MgCl_2_ and 1 M NaCl). The peak fractions were pooled, and further purified by a Superdex 200 Increase 10/300 GL column with the buffer containing 25 mM HEPES pH 7.5, 150 mM NaCl, 1 mM MgCl_2_. The peak fractions were collected and stored at -80 °C until use.

For the purification of HRAS and NRAS, the BL21(DE3) cells expressing HRAS or NRAS were resuspended in lysis buffer (25 mM Tris-HCl pH 8.0, 150 mM NaCl, 1 mM MgCl_2_) supplemented with PMSF, and disrupted by sonication on ice. The lysed cells were centrifuged with a speed of 12,000 rpm at 4 °C for 1 h, then the supernatant was incubated with TALON resin for 2 h at 4 °C, and the resin was washed with 15 column volumes of the lysis buffer, followed by 5 column volumes of wash buffer (25 mM Tris-HCl pH8.0, 500 mM NaCl, 1 mM MgCl_2_ and 5 mM imidazole). The protein was eluted with 8-10 column volumes of elution buffer (25 mM Tris-HCl pH 8.0, 150 mM NaCl, 1 mM MgCl_2_, 250 mM imidazole and 10% glycerol). The eluate was supplemented with 5 mM DTT and 0.1 mM guanosine diphosphate (GDP), diluted using buffer A (25 mM Tris-HCl pH 8.0, 1 mM MgCl_2_), then loaded onto a SOURCE 15Q (Cytiva) column and eluted by a linear gradient from 100% buffer A to 60% buffer B (25 mM Tris-HCl pH 8.0, 1 mM MgCl_2_ and 1 M NaCl). The peak fractions were pooled, supplemented with 5 mM DTT and 0.1 mM GDP, and further purified by a Superdex 200 Increase 10/300 GL column with the buffer containing 25 mM HEPES pH 7.5, 150 mM NaCl, 1 mM MgCl_2_. The peak fractions were collected and stored at -80 °C until use.

Measurement of the palmitoyltransferase activity using NBD-palmitoyl-CoA

For acyltransferase assay, {N-[(7-nitro-2-1,3-benzoxadiazol-4-yl)-methyl]amino} palmitoyl-CoA (NBD-palmitoyl-CoA), a fluorescent derivative of palmitoyl-CoA was used as a substitute of palmitoyl-CoA. In each reaction, 0.3 µM yeast or human palmitoyltransferase was first incubated with 25 µM the corresponding RAS protein in the reaction buffer (0.2 µM DTT, 1 mM EDTA and 0.02% DDM) at room temperature. For DHHC-GCP16 complexes, 0.004% CHS was added in the reaction buffer. The reaction was initiated by adding NBD-palmitoyl-CoA to the acyltransferase-RAS mixture to a final concentration of 1 µM, and quenched after 10 min (yeast) or 20 min (human) using a non-reducing SDS loading buffer. The samples were analyzed by SDS-PAGE gels. The SDS-PAGE gel was first imaged using Amersham Imager 680 (Cytiva) with a Cy2 filter (Excitation: 460 nm) (the upper panel), and then visualized by Coomassie blue staining (the lower panel).

Measurement of the autoacylation activity using a couple-enzyme assay

The couple-enzyme assay was done by following protocols developed by previous studies (Hamel et al., 2014; Rana *et al.*, 2018). The reactions were performed at 30 °C in 20 µl reaction buffer containing 50 mM sodium phosphate (pH 6.8), 1 mM EDTA, 1mM DTT, 0.02% DDM and 0.004% CHS. The reaction components included 0.25 mM oxidized nicotinamide adenine dinucleotide (NAD^+^), 0.2 mM thiamine pyrophosphate (TPP), 2 mM α-ketoglutaric acid, 3.2 mU α-KDH, 300 nM DHHC enzyme and 150 µM acyl-CoA (Avanti #870710P, #870714P, #870718P, #870720P, #870722P, Sigma #P9716). All components except acyl-CoA were incubated in the reaction buffer, then the reaction was initiated by adding acyl-CoA to a final concentration of 150 µM. The fluorescence signals were immediately measured in a microplate reader (Spark) with an excitation wavelength of 340 nm and an emission wavelength of 465 nm. The data collected in the first 10 minutes were used to calculate the reaction rates. When adding inhibitor, 100 µM 2-bromopalmitate (2-BP) was incubated with DHHC enzymes at room temperature for 1 hour before being mixed with the other components.

Cryo-EM data acquisition

Four µL of the purified sample at 8 mg/mL (Erf2-Erf4 complex) or 6-9.5 mg/mL (DHHC9-GCP16 complex) was applied onto a freshly glow-discharged holey carbon grid (Quantifoil, Au 300 mesh, R1.2/1.3). Excess sample was blotted for 3.5 s with a blot force of 0 and was vitrified by plunging into liquid ethane using a Vitrobot Mark IV (Thermo Fischer Scientific) at 8 °C and 100% humidity. Cryo-EM imaging was performed on a Titan Krios equipped with a Gatan K2 Summit direct electron detector. The microscope was operated at 300 kV accelerating voltage.

Cryo-EM image processing

A flowchart of the data processing process of *Saccharomyces cerevisiae* sample is illustrated in Figure S2. Movie stacks were motion-corrected with 2-fold binning by Motioncor2 (Zheng et al., 2017), and the resulting 15,626 dose-weighted micrographs with 1.087 Å pixel size were imported for patch-based CTF estimation by cryoSPARC v3 (Punjani et al., 2017). All the following steps were performed in cryoSPARC v3 unless otherwise stated. Through first round of particle picking and reference-free 2D classification, 514,743 high quality particles with box size of 192 pixels were selected to reconstruct an initial model (ab-initio reconstruction job). To remove different trash protein particles in 3D classification, we generated four bad references based on the initial model by “Volume Eraser” in Chimera (Pettersen et al., 2004), which represented outer membrane region only, micelle only, transmembrane region only and without transmembrane helices maps. One round of 5-classes multi-reference 3D classification (heterogeneous refinement job) was performed, yielding 195,041 particles. However, the transmembrane region density of the good class was still noisy. To achieve a better balance between signal and noise of transmembrane region, we applied a strategy called “noise reduction” to reduce the micelle noise and recover the weak signal of transmembrane helices. Briefly, a noise map of transmembrane micelle was generated by erasing signal of proteins in Chimera at first. Then, a weaker noise map was generated based on the noise map by multiplying 0.5 in Relion 3.0 (relion_image_handler –multiply_constant) (Zivanov et al., 2018). Finally, the noise-reweighted map was generated by the origin map subtracting the weaker noise map (relion_image_handler --subtract). Subsequently, five rounds of multi-reference 3D classification were performed with three references (the origin map, the noise-reweighted map and the noise map). After each round of 3D classification, the particles of the first two classes were subjected to the next round. A non-uniform refinement (Punjani et al., 2020) was performed with 123,143 particles, resulting in a 3.7 Å map (FSC=0.143). To further improve the resolution, a new round of template picking was performed, whose templates were generated by re-projection of the 3.7 Å map (template creation job), yielding about 9.57 million particles. After two rounds of 2D classification, nearly 2.87 million particles were selected to do subsequent 3D classifications and refinements. Similarly, one round of 5-classes multi-reference 3D classification and five rounds of 3-classes multi-reference (noise-reduction) 3D classification were performed, yielding 716,030 particles. The particles were then subjected to more five rounds of multi-reference 3D classification with three references generated by different lowpass filters (60 Å, 30 Å, and 10 Å). And a non-uniform refinement was performed with 556,792 particles, resulting a 3.4 Å map (FSC=0.143). All the map figures were generated using Chimera or ChimeraX (Goddard et al., 2018).

A flowchart of the data processing process of *Homo sapiens* sample is illustrated in Figure S3. A similar procedure was applied. 21,926 dose-weighted micrographs with 0.8389 Å pixel size were used. After patch-based CTF estimation, Particle picking and one round of 2D classification were performed, yielding 281,430 particles with box size of 256. Subsequently, ab initio reconstruction and non-uniform refinement were performed, resulting in a 5.5 Å map (FSC = 0.143). Then, six rounds of 3-classes multi-reference (noise-reduction) 3D classification were performed, yielding 129,114 particles. These particles were subjected to another round of multi-reference 3D classification. A non-uniform refinement with 124,470 particles was performed, resulting in a 3.9 Å map (FSC=0.143). One round of skip alignment 3D classification (relion_refine_mpi –skip_align, K=5, T=8) was performed, yielding 69,051 particles. And a local refinement was performed, resulting in a 3.7 Å map (FSC=0.143). To further improve the resolution, the strategy mentioned above was applied again. Nearly 17.47 million particles were picked. About 1.98 million particles remained after three rounds of 2D classification. Five rounds of 3-classes multi-reference (noise-reduction) 3D classification were performed, yielding 639,235 particles. Then, five more rounds of 3-classes multi-reference (low-pass gradient) 3D classification were performed, yielding 338,978 particles. Subsequently, a non-uniform refinement and one round of skip alignment 3D classification (K=5, T=4), yielding 236,228 particles. Finally, a local refinement resulted in a 3.4 Å map (FSC=0.143). All the map figures were generated using Chimera or ChimeraX.

Model building and refinement

The model of Yeast Erf2-Erf4 complex and human DHHC9-GCP16 complex were both firstly generated by AlphaFold2 Multimer (Jumper et al., 2021). The structures were docked into the density map and manually adjusted in COOT (Emsley and Cowtan, 2004). The models were refined against the final map by PHENIX (Adams et al., 2010) in real space (phenix.real_space_refine). The resolution of model versus map was determined by Fourior Shell Correlation at FSC=0.5. Statistics of map reconstruction and modelling summary can be found in table S1.

Inductively Coupled Plasma Mass Spectrometry (ICP-MS)

Element analysis was carried out using ICP-MS (Thermo Fisher Scientific, iCAP RQ) controlled with the Qtegra software package and equipped with an ASX-560 autosampler (Teledyne CETAC Technologies). Internal standard was consisted of 10 ng/mL of a mixed element solution containing ^7^Li, ^45^Sc, ^73^Ge, ^89^Y, ^115^In, ^159^Tb, ^209^Bi (General Research Institute for Nonferrous Metals, China) and added manually. Quantitative calibration and check standards were made using Zn standard (Macklin, 1000 μg/mL), Ce standard (AccuStandard, 1000 μg/mL), a mixed standard containing V, Cr, Co, Ni, Cu, Cd, Ce, Pb (General Research Institute for Nonferrous Metals, China, 20 μg/mL) and Mg, K, Fe (General Research Institute for Nonferrous Metals, China, 1000 μg/mL). The DHHC9-GCP16 complex (46.0 µM) were diluted 10 times to a concentration of 4.6 µM with a buffer of 50 mM Tris-HCl pH 7.5, 125 mM NaCl, 0.5 mM DTT, 0.02% DDM and 0.004% CHS. Both the sample and control group were then diluted 10 times with 0.02 % DDM (w/v) and 0.004% CHS (w/v) for detection. Each sample was acquired using 3 main runs. The isotopes selected for analysis were ^24^Mg, ^39^K, ^51^V, ^52^Cr, ^57^Fe, ^59^Co, ^60^Ni, ^63^Cu, ^66^Zn, ^111^Cd, ^140^Ce, ^208^Pb and ^45^Sc (chosen as an internal standard for data interpolation and machine stability).

Supplemental Figures

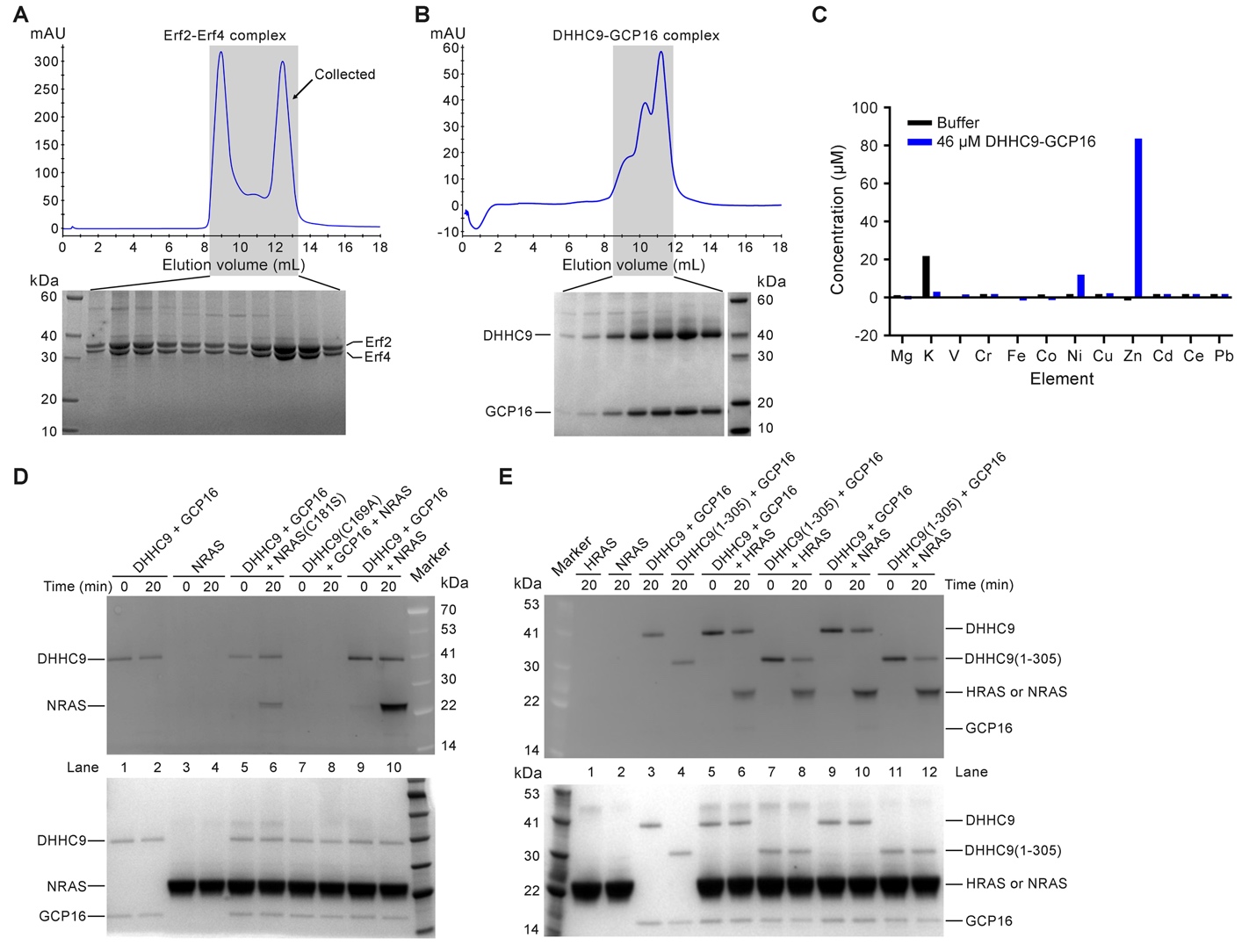

Figure S1. Purification and characterization of the Erf2-Erf4 complex and the DHHC9-GCP16 complex.

(A and B) The Erf2-Erf4 complex (A) and the DHHC9-GCP16 complex (B) purified by affinity columns were further purified by size exclusion chromatography, then the peak fractions were analyzed by SDS-PAGE followed by Coomassie blue staining.

(C) Quantification of metal ions in the purified DHHC9-GCP16 complex. The metal ions in the purified DHHC9-GCP16 complex and that in the buffer control were analyzed using ICP-MS.

(D) Palmitoylation of NRAS catalyzed by the DHHC9-GCP16 complex. The wild-type or the catalytic-dead mutant (C169A) of DHHC9 in complex with GCP16 was incubated with wild-type NRAS or its C181S mutant and NBD-palmitoyl-CoA. At the indicated time points, the reactions were quenched with non-reducing SDS loading buffer and analyzed by SDS-PAGE. The SDS-PAGE gel was first imaged using Amersham Imager 680 (Cytiva) with a Cy2 filter (Excitation: 460 nm) (the upper panel), and then visualized by Coomassie blue staining (the lower panel).

(E) The C-terminal region of DHHC9 has little effect on the palmitoyltransferase activity of the DHHC9-GCP16 complex. The palmitoyltransferase activities of the full-length and the C-terminal truncated form (residues 1-305) of DHHC9 in complex with GCP16 were measured using HRAS and NRAS as the protein substrates and NBD-palmitoyl-CoA as the palmitoyl group donor.

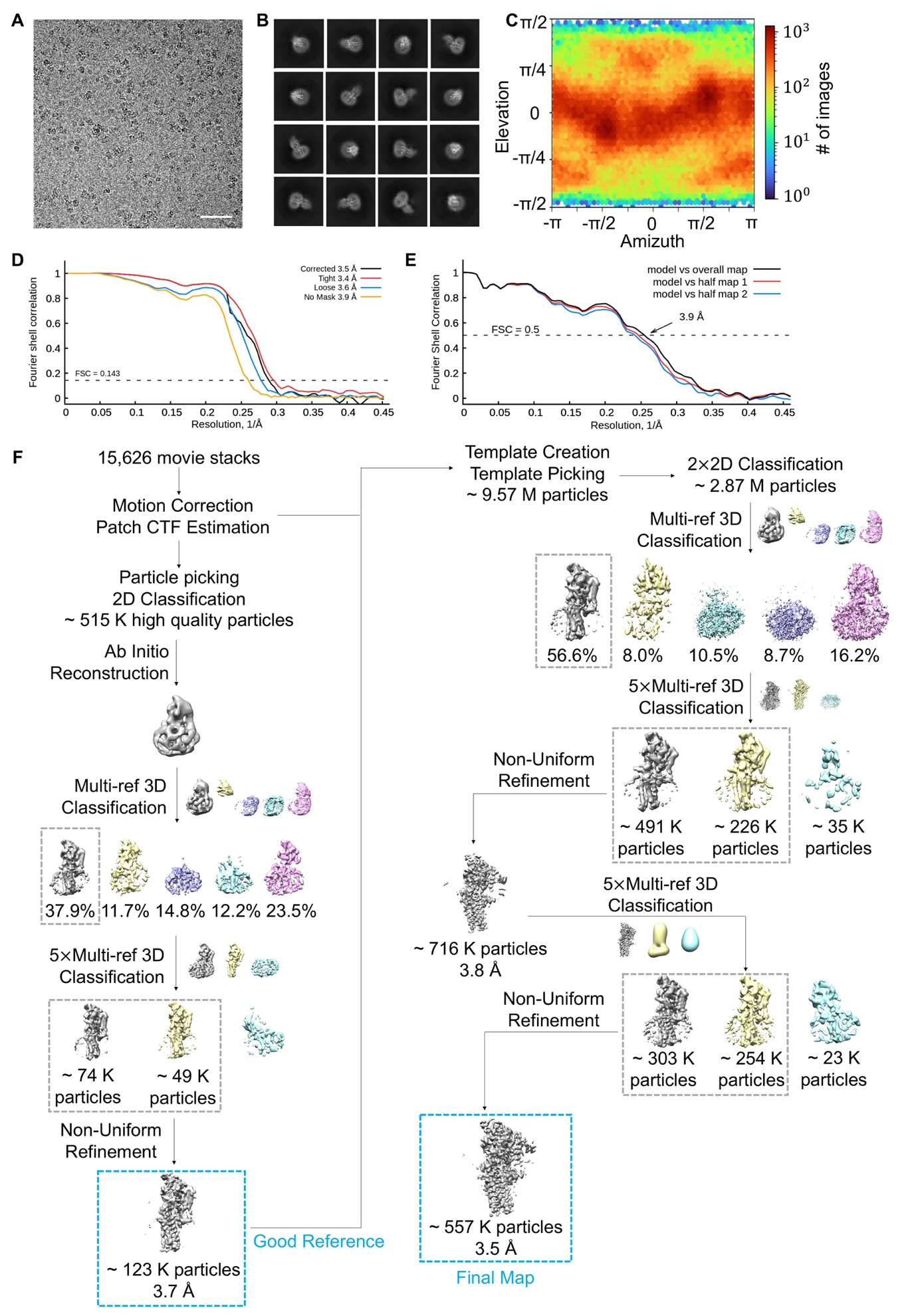

**Figure S2. Analysis of the cryo-EM data of the Erf2-Erf4 complex.**

(A) Representative cryo-micrograph. Scale bar: 50 nm.

(B) Representative two-dimensional class averages. Box size: 209 Å; Circle mask: 177 Å.

(C) Angular distribution of the particles of the final map reconstruction generated by cryoSPARC. (D) Gold standard Fourier shell correlation (FSC) curves for the 3D reconstructions generated by cryoSPARC. The curves representing the corrected (random phase initialized), tight mask, loose mask and no mask are colored black, red, blue and orange respectively.

(E) Validation of the final structure model. The curves indicate the FSC of the model versus the overall map/ half map 1/ half map 2 (FSC=0.5).

(F) A flowchart of EM data processing. Please refer to “Cryo-EM image processing” section in Methods for details.

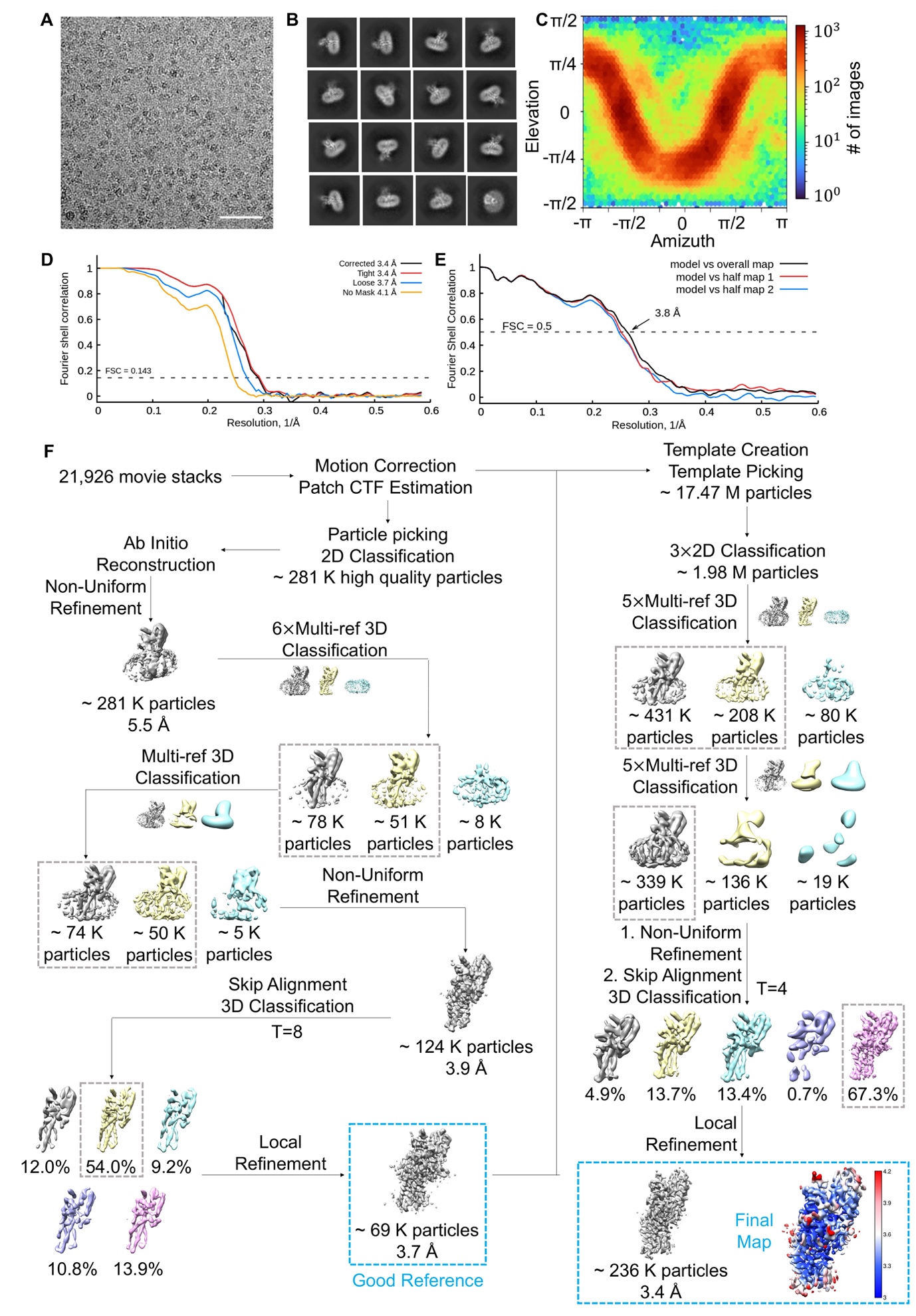

**Figure S3. Analysis of the cryo-EM data of the DHHC9-GCP16 complex.**

(A) Representative cryo-micrograph. Scale bar: 50 nm.

(B) Representative two-dimensional class averages. Box size: 215 Å; Circle mask: 183 Å.

(C) Angular distribution of the particles of the final map reconstruction generated by cryoSPARC.

(D) Gold standard Fourier shell correlation (FSC) curves for the 3D reconstructions generated by cryoSPARC. The curves representing the corrected (random phase initialized), tight mask, loose mask and no mask are colored black, red, blue and orange respectively.

(E) Validation of the final structure model. The curves indicate the FSC of the model versus the overall map/ half map 1/ half map 2 (FSC=0.5).

(F) A flowchart of EM data processing. Please refer to “Cryo-EM image processing” section in Methods for details.

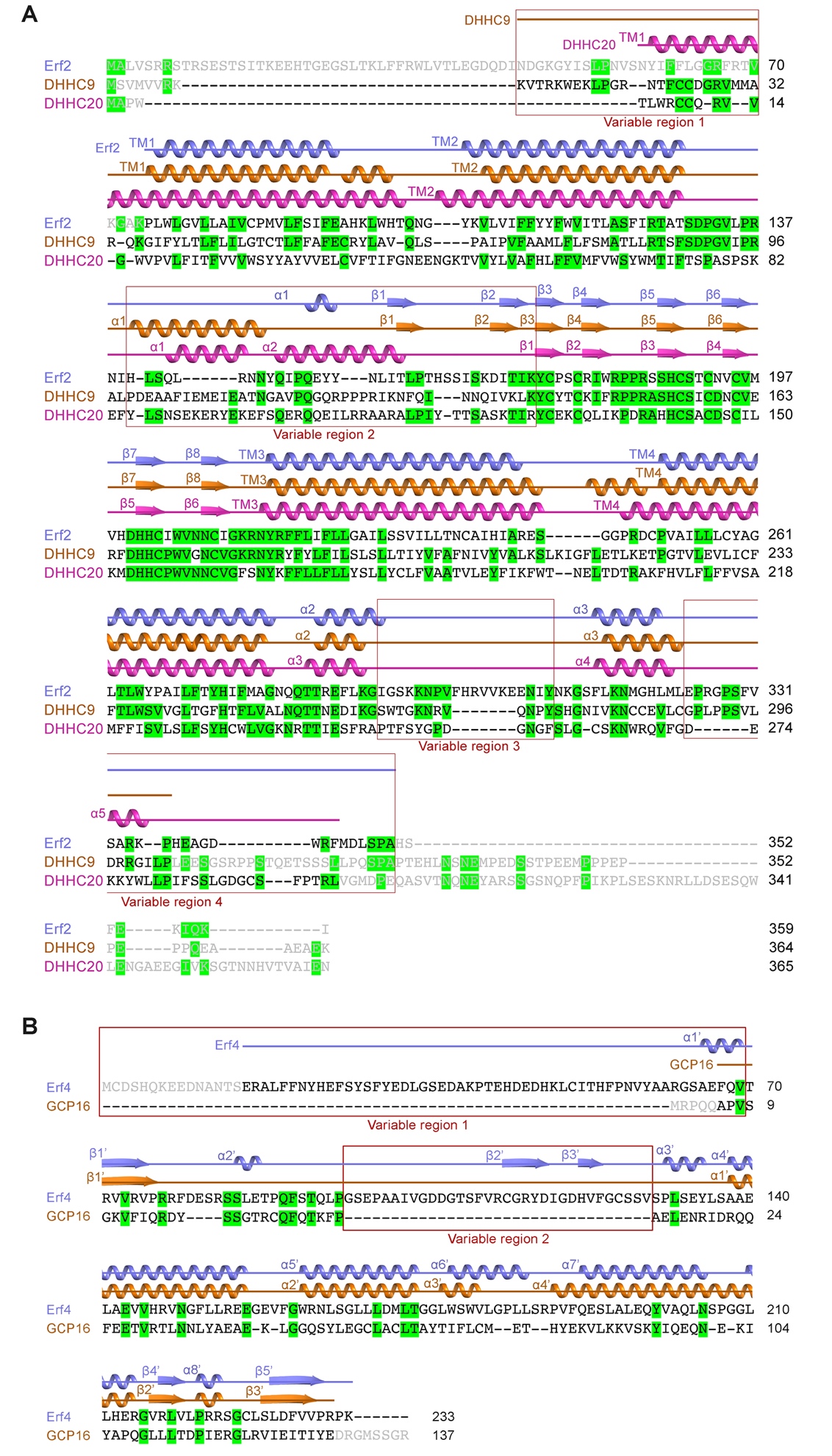

Figure S4. Structure-based protein sequence alignment.

(A) Structure-based alignment of the protein sequences of Erf2, DHHC9 and DHHC20. The protein sequences of Erf2 (UniProt ID: Q06551), DHHC9 (UniProt ID: Q9Y397) and DHHC20 (UniProt ID: Q5W0Z9) were first aligned using SnapGene (Version 6.1.2) and then the alignment was adjusted based on the cryo-EM structures of the Erf2-Erf4 complex (our study) and the DHHC9-GCP16 complex (our study) and the crystal structure of DHHC20 (PDB code: 6BMN). The residues that invisible in the structures are indicated with gray font color. The residues that are conserved in any two of the three acyltransferases are highlighted with green color. The secondary structures of Erf2, DHHC9 and DHHC20 are colored slate, orange, and magenta, respectively.

(B) Structure-based alignment of the protein sequences of Erf4 and GCP16. The protein sequences of Erf4 (UniProt ID: P41912) and GCP16 (UniProt ID: Q7Z5G4) were first aligned using SnapGene (Version 6.1.2) and then the alignment was adjusted based on the cryo-EM structures of the Erf2-Erf4 complex (our study) and the DHHC9-GCP16 complex (our study). The residues that invisible in the structures are indicated with gray font color. The residues that are conserved in any two of the three acyltransferases are highlighted with green color. The secondary structures of Erf4 and GCP16 are colored slate and orange, respectively.

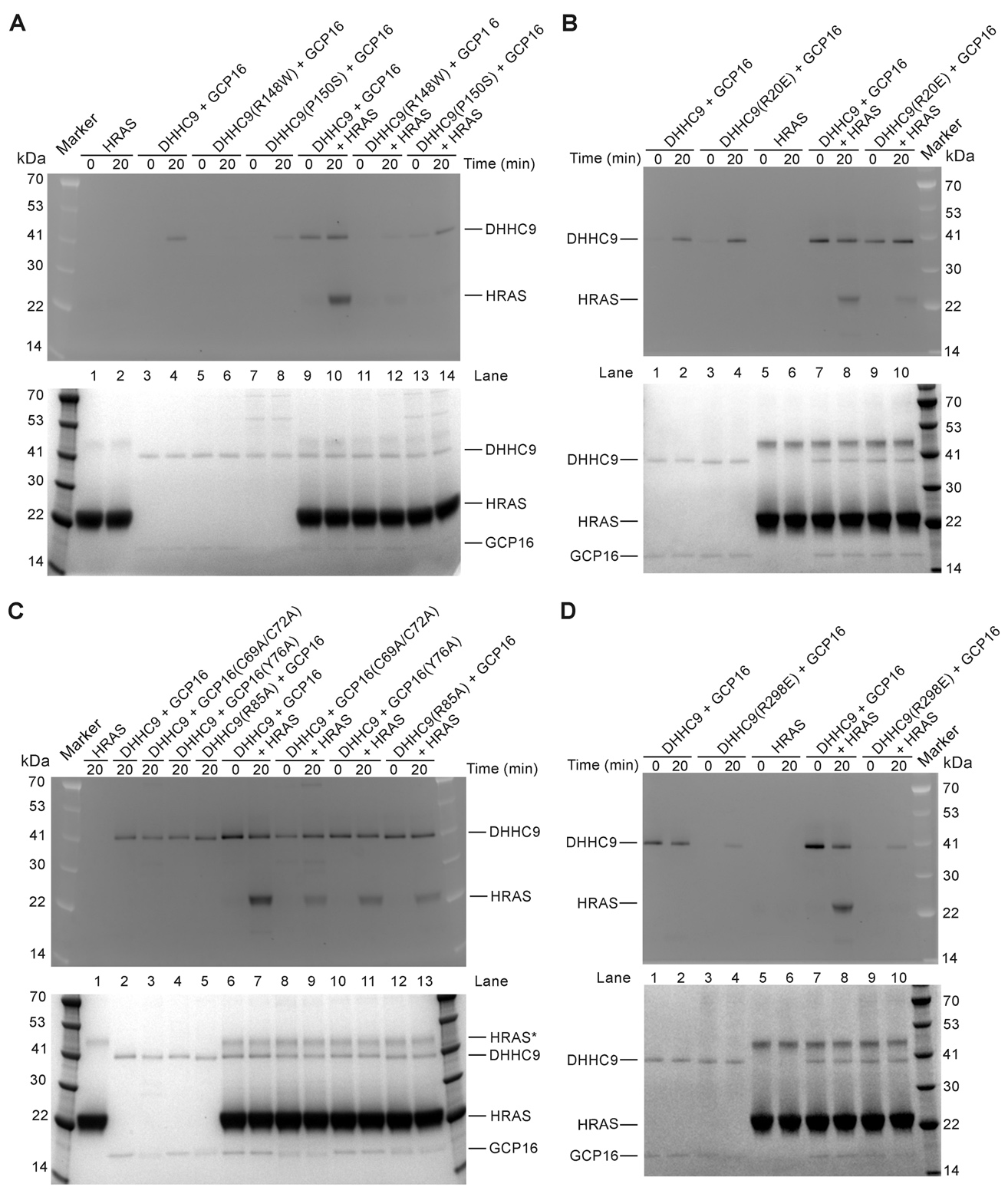

**Figure S5. Point mutations decreased the palmitoyltransferase activity of the DHHC9-GCP16 complex.**

(A) Effect of disease-causing mutations R148W and P150S in DHHC9 on the acyltransferase activity of the DHHC9-GCP16 complex.

(B to D) Effect of point mutations in DHHC9 and GCP16 on the acyltransferase activity of the DHHC9-GCP16 complex.

The palmitoyltransferase activities of the DHHC9-GCP16 complexes were measured using HRAS as the protein substrate and NBD-palmitoyl-CoA as the palmitoyl group donor. At the indicated time points, the reactions were quenched with non-reducing SDS loading buffer and analyzed by SDS-PAGE. The SDS-PAGE gel was first imaged using Amersham Imager 680 (Cytiva) with a Cy2 filter (Excitation: 460 nm) (the upper panel), and then visualized by Coomassie blue staining (the lower panel).

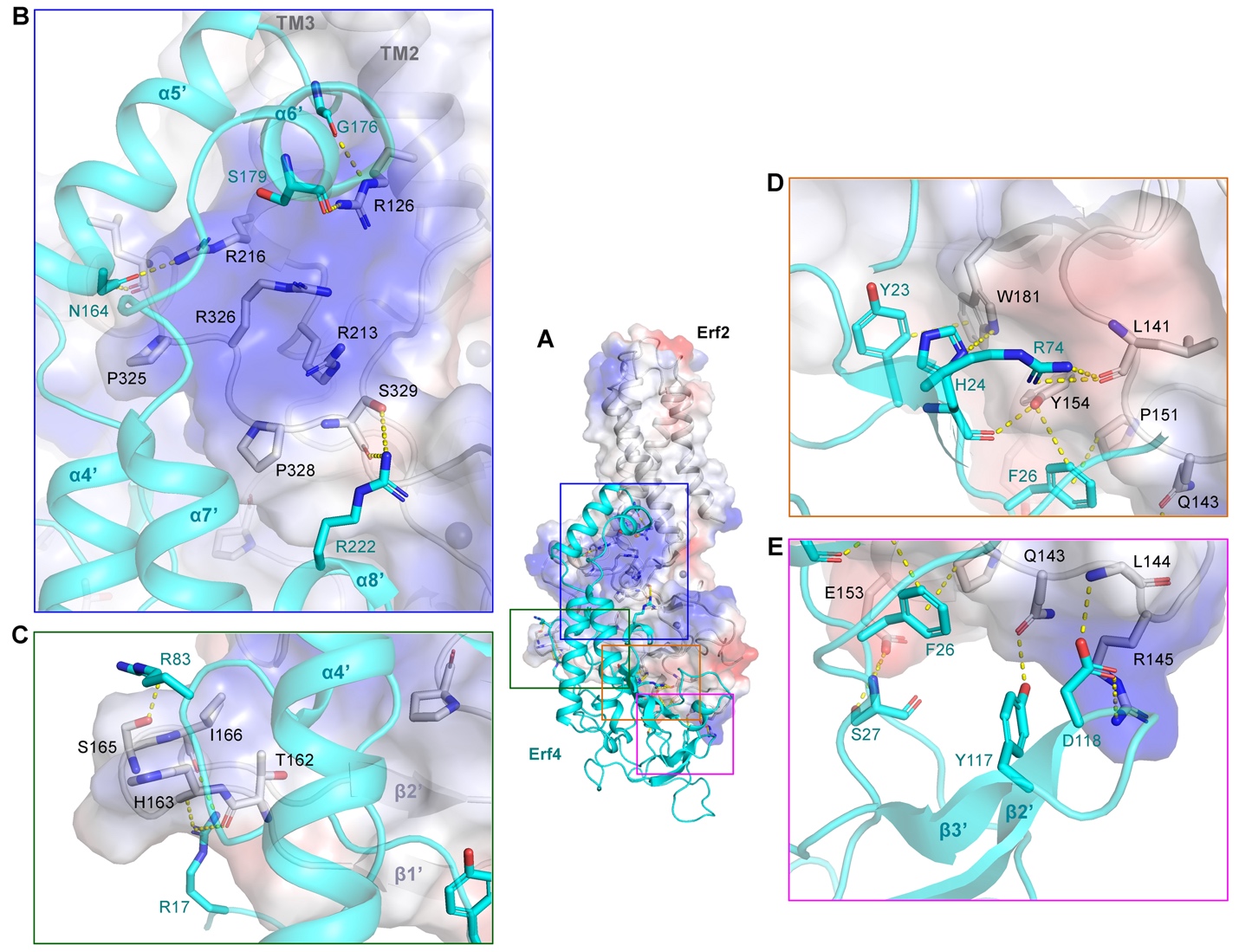

**Figure S6. Interactions between Erf2 and Erf4.**

(A) The overall structure of the Erf2-Erf4 complex.

(B) Residues R126, R216 and S329 in TM2, TM3 and the PPII helix of Erf2 interact with residues G176, S179, N164 and R222 in α4’-α8’ helices of Erf4 through hydrogen bonding. P325 and P328 in the PPII helix of Erf2 dock into to hydrophobic pockets in Erf4.

(C) Residues T162, H163, S165 and I166 in Erf2 form hydrogen bonds with residues R17 and R83 in Erf4.

(D and E) The linker between the TM2 and β1 strand of Erf2 interacts with the N-terminal region and the loop between β2’ and β3’ of Erf4 through hydrogen bonding, π–π stacking and CH-π interactions. The hydrogen bonds are represented by yellow dash lines.

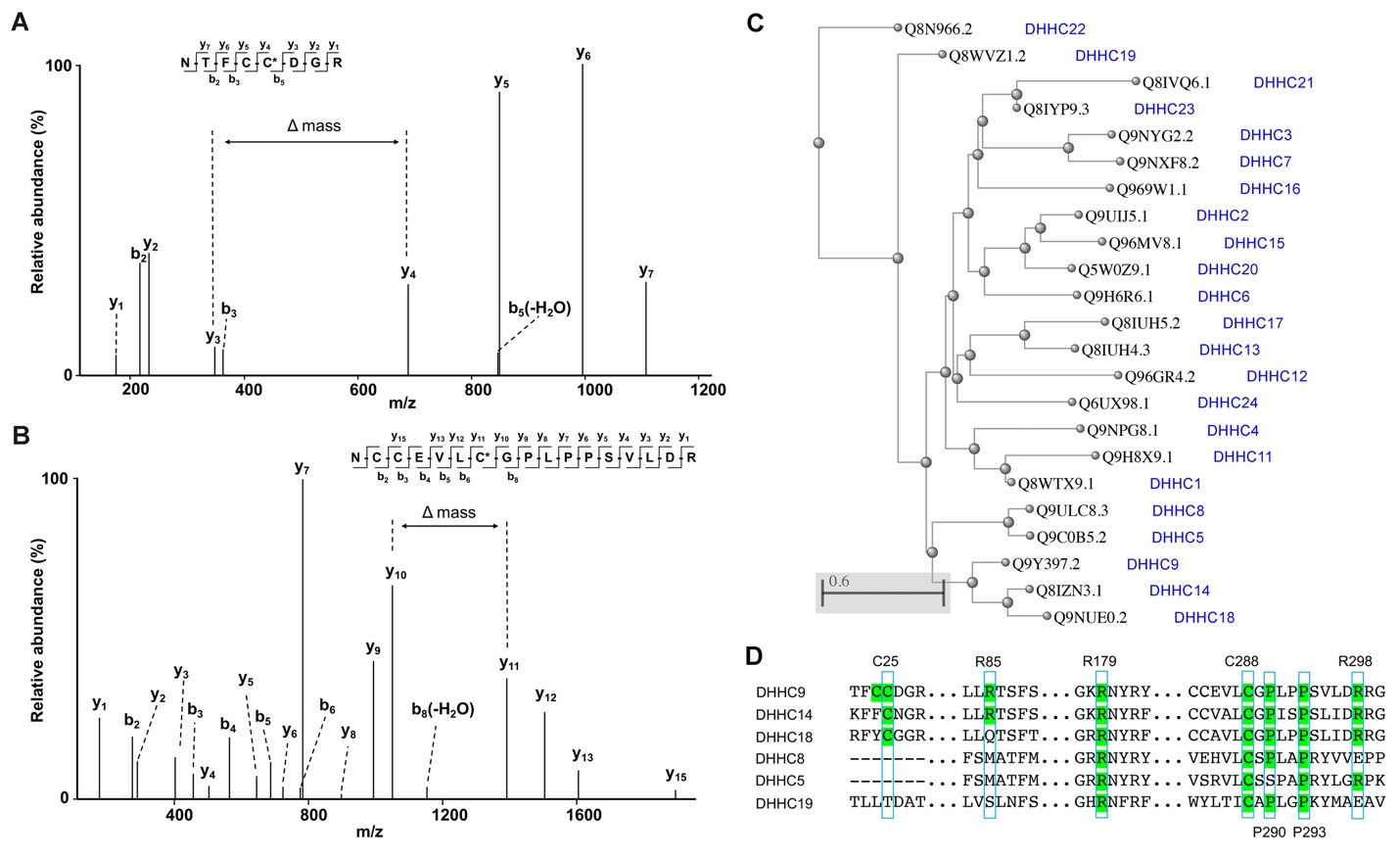

**Figure S7. Conservation of the palmitoylation sites, the phospholipid-binding residues, and the PPII helix among DHHC acyltransferases.**

(A and B) Identification of S-palmitoylation sites in DHHC9 by MS/MS. (A) The MS/MS spectra of tryptic peptides NTFCCDGR containing C24 and C25 from DHHC9. (B) The MS/MS spectra of tryptic peptides NCCEVLCGPLPPSVLDR containing C283, C284 and C288 from DHHC9. The delta mass labeled in (A) and (B) is equal to the molecular weight of palmitoylated C25 and C288, respectively.

(C) The distance tree of human DHHC acyltransferases was generated by alignment of the protein sequences using the Basic Local Alignment Search Tool (BLAST)(Johnson *et al.*, 2008). The UniProt ID and the corresponding name of each DHHC acyltransferase are showed.

(D) Alignment of the protein sequences of DHHC9 with DHHCs that share highest homology with DHHC9 using BLAST. The palmitoylated cysteine residues, the arginine residues that bind to phospholipid, and the two proline residues in the PPII helix in DHHC9 and the conserved residues in other DHHCs are colored green.

**Supplemental Tables**

**Table S1: Data collection and refinement statistics.**

|  | Yeast Erf2-Erf4  (EMD-34717)  (PDB: 8HFC) | Human DHHC9-GCP16  (EMD-34711)  (PDB: 8HF3) |
| --- | --- | --- |
| Data collection and processing |  |  |
| Microscope | FEI Titan Krios | FEI Titan Krios |
| Magnification | 81,000 | 105,000 |
| Voltage (kV) | 300 | 300 |
| Detector | Gatan K3 | Gatan K3 |
| Electron exposure (e^–^/Å^2^) | 50 | 50 |
| Defocus range (μm) | -0.5 ~ -3.0 | -0.5 ~ -3.0 |
| Pixel size (Å) | 1.087 | 0.8389 |
| Symmetry imposed | C1 | C1 |
| Initial particle images (no.) | 14,531,982 | 17,471,734 |
| Maps | Final Map | Final Map |
| Final particle images (no.) | 556,792 | 236,228 |
| Map resolution (Å) | 3.5 | 3.4 |
| FSC threshold | 0.143 | 0.143 |
| Map resolution range (Å) | 3.0-4.5 | 3.0-4.2 |
| Refinement |  |  |
| Initial model used (PDB code) | None | None |
| Model resolution (Å)  FSC threshold | 3.9  0.5 | 3.8  0.5 |
| Map sharpening *B* factor (Å^2^) | -186.6 | -172.6 |
| Model composition  Non-hydrogen atoms  Protein residues  Ligand  Water  Ion | 4007  498  1  0  2 | 3464  419  5  0  2 |
| R.m.s. deviations  Bond lengths (Å)  Bond angles (°) | 0.007  0.656 | 0.006  0.671 |
| Validation  MolProbity score  Clashscore  Poor rotamers (%) | 2.37  9.17  3.88 | 1.77  10.96  0.81 |
| Ramachandran plot  Favored (%)  Allowed (%)  Disallowed (%) | 93.32  6.48  0.20 | 96.63  3.37  0.00 |
